# Astrocytes expressing M_4_ muscarinic acetylcholine receptor regulate locomotion and survival in murine prion disease

**DOI:** 10.1101/2025.01.29.635510

**Authors:** Gonzalo S. Tejeda, Colin Molloy, Carrie K. Jones, Craig W. Lindsley, Graeme Milligan, Andrew B. Tobin

## Abstract

Whilst there has been much focus on neuronally expressed muscarinic acetylcholine receptors (mAChRs), less attention has been paid to their expression and role in non-neuronal brain cells. Using genetically engineered mice, we identify a previously unappreciated subpopulation of astrocytes expressing the M4 mAChR subtype dispersed across various regions that include the brainstem, hypothalamus and, most abundantly, the cerebellum. Signalling and proteomic analysis of M4 mAChR positive astrocytes from the cerebellum confirmed functional receptor expression and its role in regulating protein expression. Genetic ablation of these astrocytes in mice revealed a specific role in locomotion. Importantly, in the context of murine prion disease, we observed a significant expansion of M4 mAChR positive astrocytes, and ablation experiments established that this astrocyte subpopulation has a detrimental effect on late-stage disease. Together we identify M4 mAChR expressing astrocyte subpopulation that plays a specific role in normal neurophysiology and the progression of inflammatory neurodegenerative disease.

## Background

The muscarinic acetylcholine receptors (mAChRs) have been under intense investigation due to the critical role of this family of G protein-coupled receptors (GPCRs) in the regulation of various neuronal processes, including learning, memory, reward and seek behaviour. Among the five subtypes (M_1_-M_5_ mAChRs), M_1_ and M_4_ have emerged as particularly important in brain function and they are considered promising therapeutic targets for the treatment of several brain disorders (1, 2). However, while most research on mAChRs in the central nervous system (CNS) has focused on neuronal expression and function, much less is known about their roles in non-neuronal cells.

Some of the strongest evidence for mAChRs function in glial cells comes from studies on M_1_ mAChR in oligodendrocyte precursor cells (OPCs), where specific ablation of this receptor gene enhances remyelination in mouse models of autoimmune encephalomyelitis (3), hypoxia (4), aging (5) and Alzheimer’s disease (6). Additional clues suggest that mAChRs may regulate neuroinflammation. For example, our studies in a prion model of neurodegeneration (7), and others in mouse models of Alzheimer’s disease (8), have shown that the activation of the M_1_ mAChR by selective positive allosteric modulators (PAMs) reduced markers of astrocyte and microglial activation. However, whilst transcripts for the mAChR family have been reported in astrocytes and microglia (9, 10), including the M_4_ mAChR (11–13), little is known about the contribution that these receptors play in the regulation of glial cells.

Here, we used a transgenic mouse model expressing CRE recombinase under the control of the M_4_ mAChR gene promoter (*Chrm4*) to discover a discrete population of M_4_ mAChR-expressing astrocytes located in the brainstem, hypothalamus, and cerebellum. Signalling and proteomic studies confirmed the functional expression of this receptor subtype in astrocytes, particularly within the cerebellum. CRE-mediated deletion of M_4_ mAChR expressing astrocytes exposed the role of this subpopulation of astrocytes in locomotion. Importantly, our study unveiled the expansion of the M_4_ mAChR expressing astrocytes in parallel with the progression of prion neurodegenerative disease. Genetic ablation of these astrocytes significantly extended the lifespan of prion-diseased mice, suggesting a detrimental role in disease progression. Together our data point to a previously unappreciated subpopulation of M_4_ mAChR expressing astrocytes that play a specific role in normal neurophysiology and inflammatory neurodegenerative disease.

## Results

### Generation of a genetic mouse model to identify M_4_ mAChR expressing cells

To facilitate the study of the diversity and distribution of M_4_ mAChR expressing cells (M_4_^+^ cells) in the brain, we generated a mouse line that expresses a CRE recombinase under the control of the *Chrm4*-promoter (Chrm4-CRE-IRES-mM4, hereinafter referred to as Chrm4^CRE^) (**Fig. 1a**). In these mice, the *Chrm4* locus was modified to contain an internal ribosomal entry site (IRES) cassette followed by the CRE recombinase coding sequence down-stream of the M_4_ mAChR receptor exon 3 (**Fig. 1a**). By crossing these mice with a CRE-dependent green fluorescent protein (GFP) reporter mouse strain (R26-CAG-GFP,-MetRS*^L274G^, also called R26-MetRS*), we aimed to generate animals where the GFP acts as a marker of M_4_ mAChR expressing cells (**14) (Fig. 1a**). The resulting mouse strain (Chrm4^CRE^::MetRS*), heterozygous for both alleles, showed similar levels of M_4_ mAChR expression to that of wild-type mice as determined by radio-ligand binding experiments of striatal membranes **(Supplementary Fig. 1a)**. Examination of brain sections of Chrm4^CRE^::MetRS* animals revealed GFP-positive neurons (stained with the neuronal marker MAP2) especially abundant in the striatum, cortex, and hippocampus (**Fig. 1b**), areas where Western blots confirmed M_4_ mAChR expression **(Supplementary Fig. 1b)** consistent with previous studies (15–18). These experiments, together with the lack of GFP signal detected in the brain of the parental lines **(Supplementary Fig. 1c)**, give confidence that the reporter-GFP system acts as a surrogate marker for M_4_ mAChR expressing cells/neurons.

**Fig. 1:**
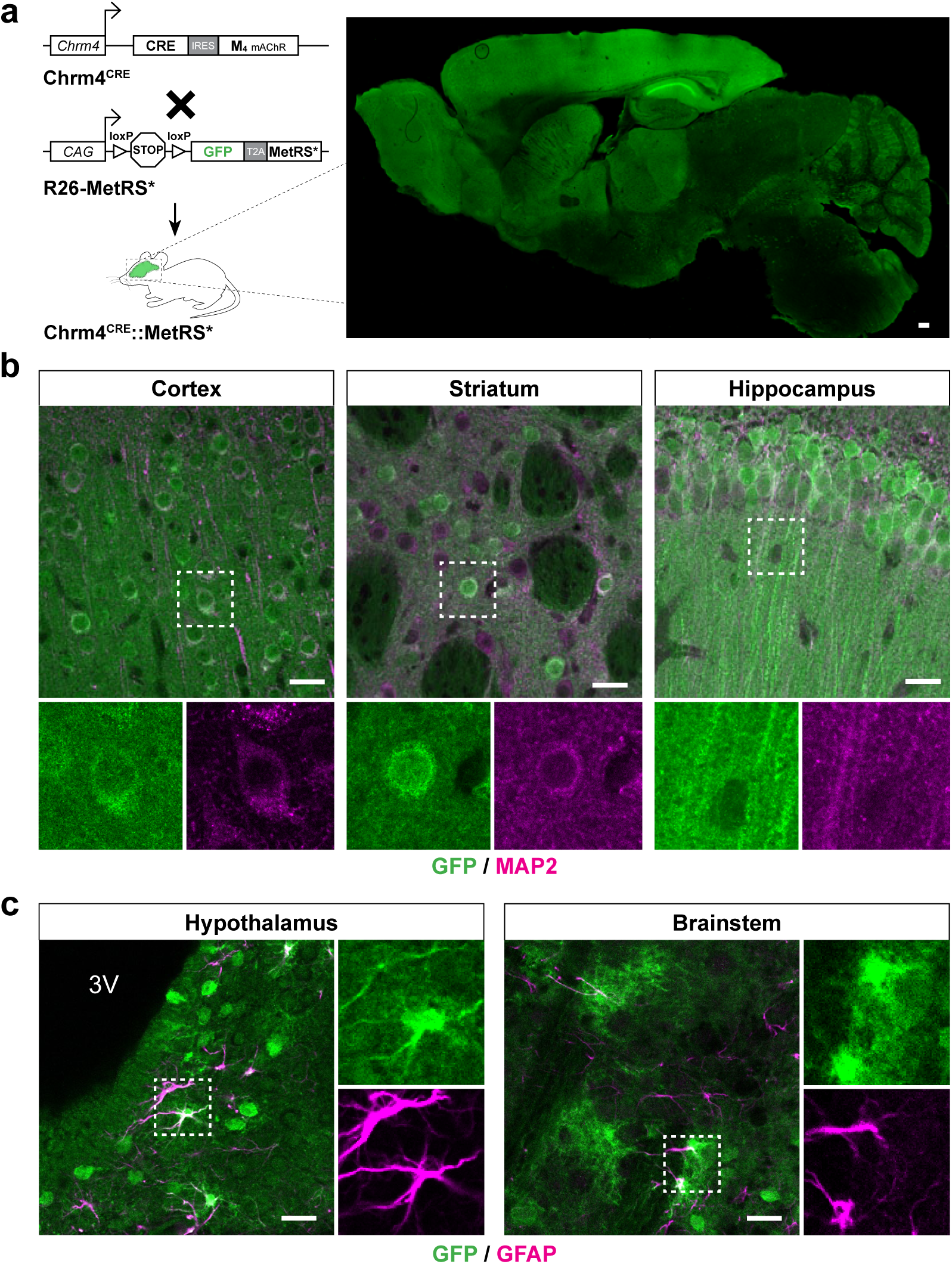
M_4_ mAChR is expressed in neuronal and disperse astrocytic populations throughout the brain. **a**, Schematic overview of the breeding between mice driving CRE recombinase from the *Chrm4* promoter (Chrm4^CRE^) with mice carrying the inducible transgene for expression of GFP and MetRS* L274G (R26-MetRS*). The distribution of M_4_ mAChR expressing cells can be detected in the resulting Chrm4^CRE^::MetRS* mice by GFP immunostaining (green) in a brain sagittal section. **b**, Representative confocal images showing co-immunostaining of GFP (green) with the neuronal marker MAP2 (magenta) in areas of the cortex, striatum and hippocampus. The white square insets indicate the enlarged areas showed in the panels below by each individual channel. **c**, Representative confocal images showing co-localisation of GFP expressing cells with the astrocyte marker GFAP (magenta) in the hypothalamus and the brainstem. Scale bar for panel **a** is 250 µm, and for **b** and **c**, 25 µm.

### M_4_ mAChRs expression in a discrete astrocytic subpopulation

In addition to M_4_^+^ neurons, we also detected dispersed populations of M_4_^+^ stellate cells in the midbrain, hypothalamus, and the pons (**Fig. 1c**). These cells did not co-localise with markers for microglia, oligodendrocytes or neurons **(Supplementary Fig. 1d)** but rather co-stained with antibody markers for glial fibrillary acidic protein (GFAP) and S100β, thereby identifying them as M_4_^+^ astrocytes (**Fig. 1c, Supplementary Fig. 1e)**.

These experiments also revealed a surprisingly diverse array of M_4_^+^ cells in the cerebellum (**Fig. 2a),** a region with low but detectable levels of M_4_ mAChR transcript **(Supplementary Fig. 2a)**. In contrast, no significant GFP staining was observed in the cerebellum of the parental mouse strains **(Supplementary Fig. 2b)**. Among these M_4_^+^ cells, we could identify a sparse population of M_4_^+^ Purkinje neurons, labelled with the calcium-binding protein calbindin D-28k (**Fig. 2b**). Most strikingly, however, was the abundance of cerebellum M_4_^+^ cells that co-stained with astrocytic markers (GFAP, NDRG2, S100β) (**Fig. 2c, Supplementary Fig. 2c)**. The location and morphology of these cells indicated that they correspond to Bergmann glia in the Purkinje layer, some velate astrocytes in the granule cell layer, dispersed fibrous astrocytes surrounding the white matter and a large population of M_4_^+^ astrocytes within the cerebellar nuclei (**Fig. 2c).**

**Fig. 2:**
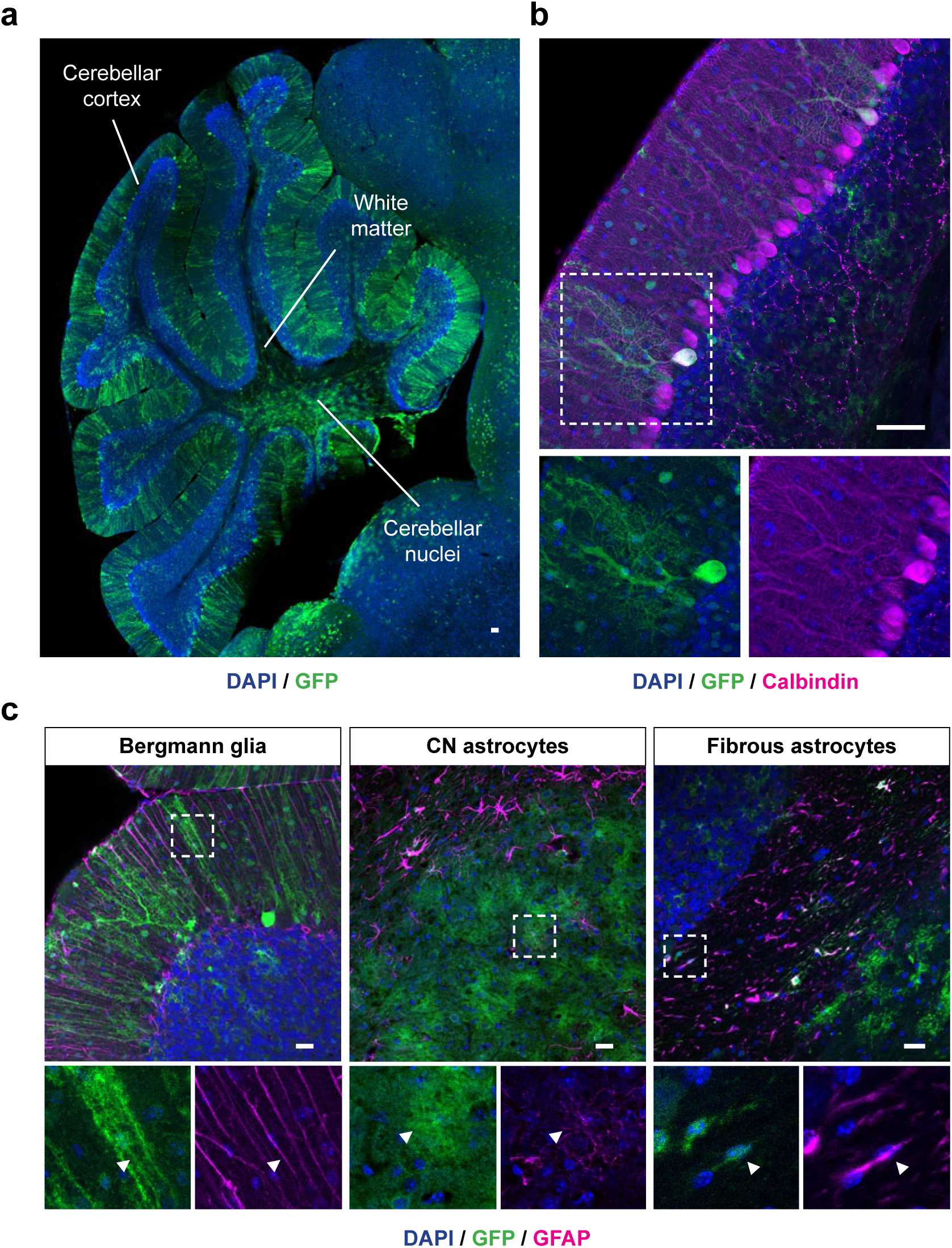
The cerebellum contains a diverse array of cells expressing M_4_ mAChR. **a**, Distribution of M_4_ mAChR expressing cells labelled with GFP staining in a sagittal section of the cerebellum of Chrm4^CRE^::MetRS* mice. **b**, Co-immunostaining of GFP-labelled M_4_^+^ cells with Calbindin in the Purkinje layer of the cerebellum of Chrm4^CRE^::MetRS* mice. **c**, Immunohistochemistry analysis of different populations of GFAP positive astrocytes (magenta) co-expressing M_4_ mAChR (green) in the cerebellar cortex (left panel), cerebellar nuclei (CN, middle panel) and white matter region (right panel). Arrowheads indicate M_4_^+^ astrocytes. Scale bar for panels **a** and **b** is 250 µm, and for **c**, 25 µm.

### Signalling analysis of astrocyte M_4_ mAChRs

To determine the functional expression of M_4_ mAChRs in glial cells, primary astrocyte cultures were prepared from the cerebrum and the cerebellum of Chrm4^CRE^::MetRS* mice. In line with the observations in intact brain, M_4_^+^ astrocytes represented only a small proportion of the total astrocyte population prepared from the cerebrum (8.7 ± 2.4 %), whilst a higher proportion (30.4 ± 3.8 %) of M_4_^+^ astrocytes were evident in cerebellar cultures (**Fig. 3a, b**).

**Fig. 3:**
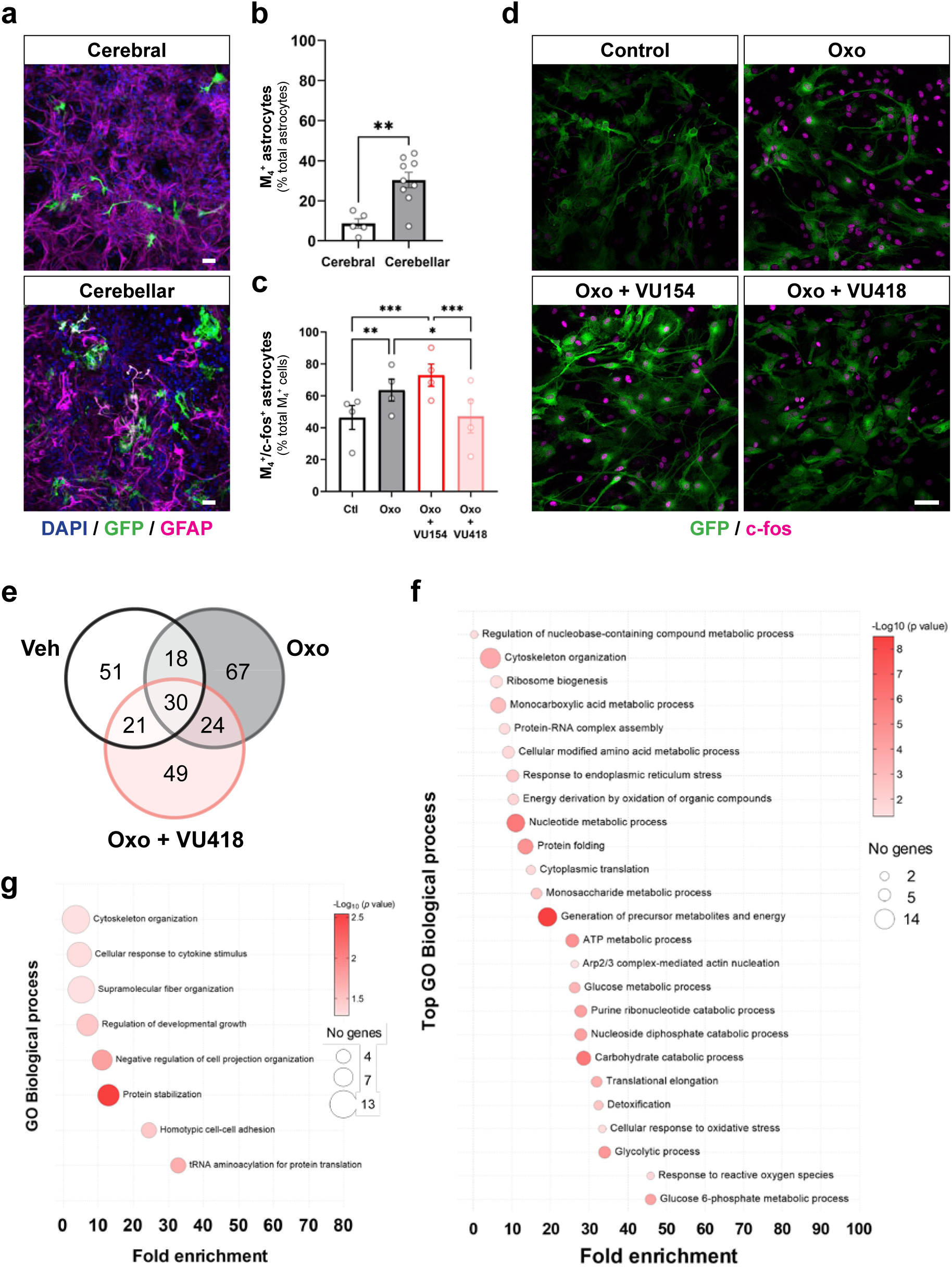
Role of M mAChR in cerebellar astrocytic cultures of Chrm4^CRE^::MetRS*. **a**, Immunostaining of cerebral or cerebellar astrocyte cultures from Chrm4^CRE^::MetRS* animals detecting M_4_ expressing cells with GFP (green) and the astrocyte marker GFAP (magenta). **b**, Quantification of the proportion of M_4_^+^ astrocytes in cerebral and cerebellar cultures relative to the total astrocytes present in the preparation (n = 5 - 9). **c**, The percentage of the population of M_4_^+^ astrocytes co-expressing c-fos was estimated in astrocyte cerebellar cultures after stimulation with Oxotremorine (Oxo, 10 µM), a combination of Oxo and the M_4_ mAChR PAM VU0467154 (VU154, 10 µM) or Oxo plus the M_4_ mAChR antagonist VU6028418 (VU418, 5 µM) for 60 min (n= 5). **d**, Representative confocal images showing M_4_ expressing astrocytes (green) co-stained with c-fos (magenta) in cerebellar cultures after treatment with Vehicle (Control), Oxo or a combination of Oxo and VU154 or Oxo with VU418, as indicated in **c**. **e**, Venn diagram showing the newly synthesised proteins (NSPs) exclusively identified from M_4_^+^ cerebellar astrocytes after treatment with ANL (4 mM) in combination with Vehicle, Oxo (10 µM) or Oxo and VU418 (5 µM). **f**, Analysis of enriched GO annotations (Fisher’s exact test performed with Panther v. 17.0) in M_4_^+^ cerebellar astrocytes after treatment with ANL in basal conditions (n = 2). The size of each circle represents the number of genes, and the colour intensity indicates the -log_10_ *p* value adjusted by Benjamini-Hochberg method assigned to the corresponding GO term. **g**, Gene ontology biological processes which are significantly enriched from the nascent proteins synthesized after activation of M mAChR in M_4_^+^ cerebellar astrocytes (NSPs exclusively found after Oxo treatment and not identified when the M_4_ antagonist is present). Graphs depict means ± SEM and statistical significance was calculated by two-tailed unpaired *t* test (**b**), or one-way ANOVA followed by Tukey’s pot hoc test (**c**) (\**p* < 0.05, \*\**p* < 0.01, \*\*\**p* < 0.01,). Scale bars, 25 µm.

To establish if these M_4_^+^ astrocytes expressed functional M_4_ mAChRs, we took advantage of previous studies that showed that muscarinic agonists can induce accumulation of the Immediate Early-Gene (IEG) c-fos in cultured astrocytes (19). Therefore, we treated cerebellar astrocytes prepared from Chrm4^CRE^::MetRS* mice with the muscarinic agonist oxotremorine (10 μM). This resulted in a significant up-regulation of c-fos expression in M_4_^+^ astrocytes (**Fig. 3c, d**). Furthermore, preincubation with VU0467154 (VU154, 10 μM), a M_4_ mAChR PAM (20), increased the number of oxotremorine-induced c-fos^+^ cells in the cerebellar astrocyte cultures, whilst pretreatment with the highly selective M_4_ mAChR antagonist VU6028418 (VU418, 5 μM) (21) prevented oxotremorine-induced stimulation of c-fos in these cells (**Fig. 3c, d**). Additionally, we probed the phosphorylation of the extracellular signal-regulated kinases 1/2 (pERK_1/2_), a signalling pathway downstream of M_4_ mAChR-Gi activation. Treatment with oxotremorine increased pERK_1/2_ levels in a concentration-dependent manner (pEC_50_ = 5.57 ± 0.21, **Supplementary Fig. 3a**) whereas the M_4_ mAChR antagonist VU418 (1 μM) shifted the curve to higher agonist concentrations (pEC_50_ = 4.61 ± 0.22, *p* = 0.002) consistent with VU418 selective antagonism of the M_4_ mAChR ERK_1/2_ response.

### Proteomic evaluation of astrocyte M_4_ mAChRs

To further characterise the functional M_4_ mAChR response in cerebellar astrocytes, we analysed changes in *de novo* synthesised proteins in response to receptor activation. In these experiments we took advantage of the fact that, in addition to the expression of GFP, the Chrm4^CRE^::MetRS* mice contain a mutated methionyl-tRNA synthetase (MetRS L274G) expressed in a CRE-dependent manner. The enlarged binding pocket of this mutant synthetase allows for the incorporation of the unnatural amino acid azidonorleucine (ANL) into nascent proteins, which can later be tagged by click chemistry for identification of labelled proteins or for biochemical purification (14). Since our system results in CRE-expression only in M_4_^+^ cells (i.e. astrocytes) then the mutated methionyl-tRNA synthetase will only be expressed in M_4_^+^ astrocytes. In this way, although our cerebellar astrocyte cultures are mixed (∼30% of the astrocytes are M_4_^+^ (**Fig. 3b**)), we could conduct proteomic analysis on proteins synthesised exclusively in M_4_^+^ astrocytes.

We first tested our system by exposing cerebellar astrocyte cultures, prepared from Chrm4^CRE^::MetRS* mice, to ANL (4 mM) followed by fixation, permeabilization and labelling ANL-incorporated proteins with the fluorescent Cy3-alkyne tag – process referred to as fluorescent non-canonical amino acid tagging (FUNCAT). This established a FUNCAT signal (Cy3) that was restricted to M_4_^+^ astrocytes (identified by GFP and S100β) **(Supplementary Fig. 3b)**.

Next, we collected lysates from ANL treated cerebellar astrocytes prepared from Chrm4^CRE^::MetRS* mice and using click chemistry incorporated a biotin-tag to ANL incorporated proteins. Western blot analysis revealed an abundance of biotinylated nascent proteins using this approach **(Supplementary Fig. 3c)**. These biotinylated proteins were purified using avidin beads and proteins identified by mass spectrometry **(Supplementary Data 1)**. These experiments detected a total of 260 enriched biotinylated proteins (**Fig. 3e**). A gene ontology (GO) analysis revealed enrichment of astrocytic markers (GFAP, vimentin, 3-phosphoglycerate dehydrogenase), confirming the specific labelling. Importantly, terms associated with metabolic enzymes, translation and cytoskeleton components were among the most significantly enriched GO groups (**Fig. 3f**). Furthermore, oxotremorine (10 **m**M) treatment induced the synthesis of 67 unique proteins that were dependent on M_4_ mAChR activation, as they were not identified in the presence of the specific M_4_ mAChR antagonist (VU418, 5 **m**M). A GO enrichment analysis showed that proteins up-regulated by M_4_ mAChR activation are involved in trafficking and protein translation (**Fig. 3g**), including the actin-binding profilin-2, the metallothionein subtypes II (MT2) and III (MT3), guanylate-binding protein 2 (GBP2), and the MAP kinase ERK_2_.

### Diptheria toxin-mediated ablation of M_4_^+^ astrocytes effects locomotion

In order to decipher the role of M_4_^+^ astrocytes, we followed a genetic strategy of targeted ablation. The mouse strain Aldh1l1-eGFP,-DTA (22), expresses eGFP in astrocytes and a CRE-dependent diphteria toxin A (DTA). Crossing these animals with the Chrm*4^CRE^* strain results in a mouse (Chrm4^CRE^:: Aldh1l1^DTA^) where M_4_^+^ astrocytes are selectively ablated (**Fig. 4a**). As M_4_ mAChR expression was observed in some hypothalamic astrocytes (**Fig. 1b**), we first evaluated the body weight of these animals over time and found no differences compared to wild type littermates (**Fig. 4b**). Also, these animals showed similar food consumption and organ weights **(Supplementary Fig. 4a, b)**. Previous studies have indicated that alterations in cerebellar astrocytes, specifically Bergmann glia, led to cerebellar atrophy and motor impairment (23). However, we did not observe any abnormalities in the morphology or the overall glial composition of the cerebellum of the Chrm*4^CRE^*:: Aldh1l1^DTA^ animals when compared to the cerebellum of the parental line **(Supplementary Fig. 4c)**. Furthermore, the Chrm4^CRE^:: Aldh1l1^DTA^ mice did not show ataxia or any deficits in motor coordination as determined by the hindlimb clasping, ledge and rotarod tests (**Fig. 4c**). In contrast, exploratory behaviour evaluated in an open field test showed that Chrm4^CRE^:: Aldh1l1^DTA^ mice exhibited significantly increased time in the centre of the arena compared to littermate controls (74.1 ± 6.7 and 51.4 ± 2.6 s, respectively, *p* < 0.01, **Fig. 4d**), while the overall locomotion was reduced (24.1 ± 1.5 m travelled in the arena versus 30.6 ± 1.7 m for control mice) and the time spent immobile (231.0 ± 13.7 s) was greater than wild type littermates (164.7 ± 14.4 s) (**Fig. 4e, Supplementary Fig. 4d)**. These results could be interpreted as deficits in anxiety-like behaviour, but a further evaluation in an elevated plus maze test excluded this possibility, as Chrm4^CRE^:: Aldh1l1^DTA^ mice performed similarly to wild type littermates in this paradigm (**Fig. 4e, Supplementary Fig. 4e)**.

**Fig. 4:**
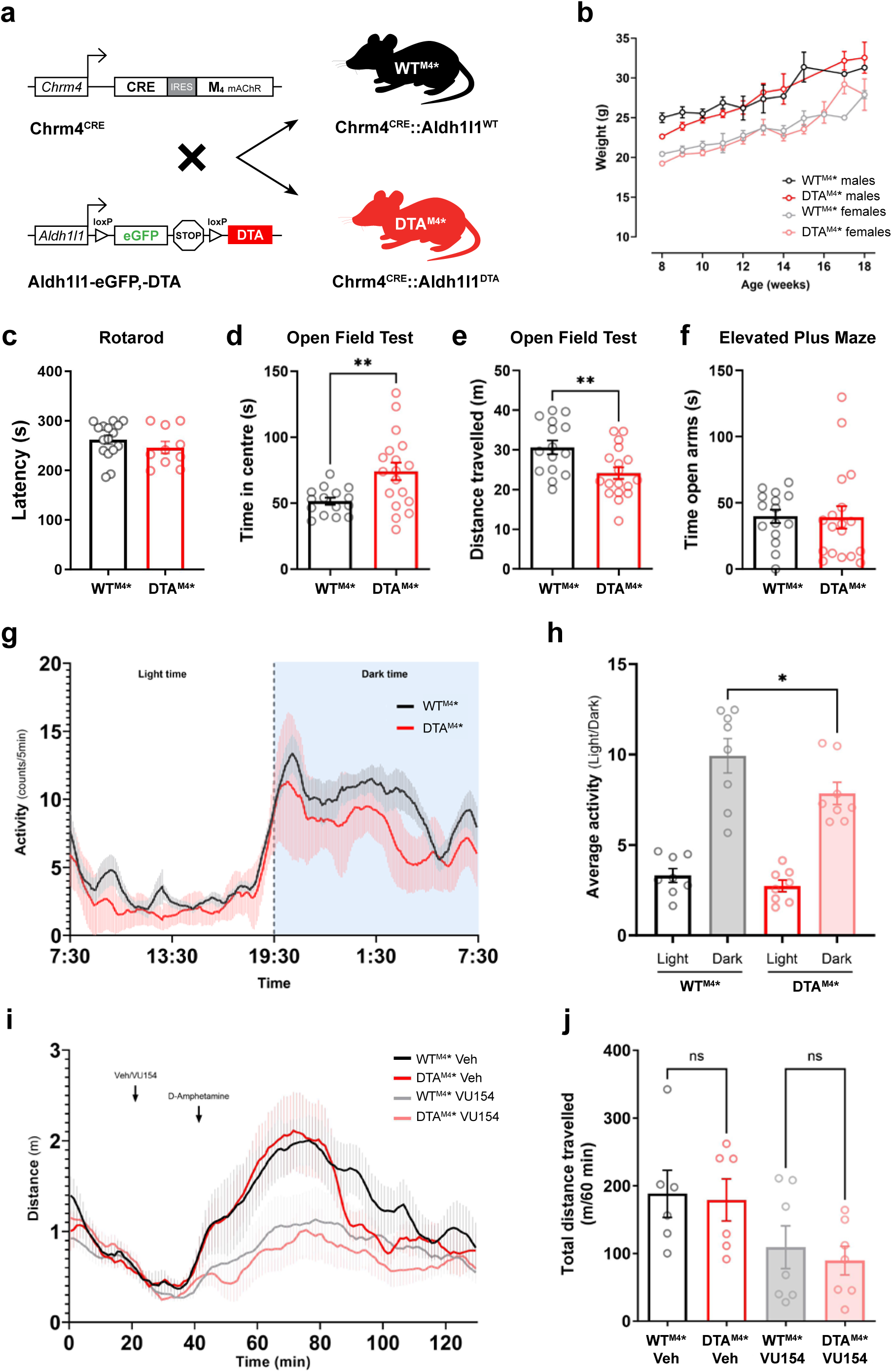
Selective ablation of M_4_^+^ astrocytes leads to deficits in locomotor activity. **a**, Depiction showing the breeding overview between the Chrm4^CRE^ animals with the hemizygous Aldh1l1-eGFP,-DTA mice, that express eGFP under the astrocyte promoter *Aldh1l1* and an inducible diphteria toxin A when Cre recombinase is present. This mating results in Chrm4^CRE^:: Aldh1l1^DTA^ mice (DTA^M4^*) with ablated astrocytes that express the M mAChR, and the control littermates Chrm4^CRE^:: Aldh1l1^WT^ (WT^M4^*), wild type for the floxed gene. **b**, Average animal weight per group over time (n = 6-10). **c**, Comparison of the performance on the accelerated rotarod test between DTA^M4^* and WT^M4^* mice during 4 consecutive weeks (n = 10). **d**, Activity of mice in and open field test measuring the time spent in the central area of the apparatus (n = 15-18). **e**, Total distance travelled over the whole area of the open field arena. **f**, Analysis of the anxiety-like behaviour of DTA^M4^* and WT^M4^* mice measuring the time spent in the open arms of an elevated plus maze (n = 15-18). **g**, Locomotor activity in DTA^M4^* animals compared to WT^M4^* littermates measured by telemetry. The graph depicts the profile of activity counts averaged over 3 days of recording and plotted every five min. Mice experience a 12/12 h light/dark cycle, with the dark cycle shown in the shaded area (n = 8). **h**, Comparison of the average telemetry activity counts during the 12-h light and dark phases between DTA^M4^* and WT^M4^* mice. **i**, Effect of VU0467154 (VU154, 10 mg/kg) or vehicle (Veh) to counteract the amphetamine-induced hyperlocomotion (2 mg/kg) in DTA^M4^* and WT^M4^* mice. **j**, Total distance travelled during the 60 min period following D-amphetamine administration. (n = 6-7). Data are means ± SEM and statistical significance was calculated by two-tailed unpaired *t* test or one-way ANOVA followed by Bonferroni’s pot hoc test when analysis involved more than two groups (\**p* < 0.05, \*\**p* < 0.01).

These data suggest a potential role for M_4_^+^ astrocytes in locomotion which was tested further by implant telemetry. Both Chrm4^CRE^:: Aldh1l1^DTA^ and wild type littermate mice had peaks of activity at similar times and of similar duration, excluding the possibility of disturbed sleep (**Fig. 4g**). Moreover, fluctuations in body temperature followed a similar pattern, with no differences observed between the two groups of animals **(Supplementary Fig. 4f)**. Interestingly, when considering the overall activity in each phase, it was evident that Chrm4^CRE^:: Aldh1l1^DTA^ mice moved less intensely during the dark phase (7.85 ± 0.61 counts) than their control littermates (9.93 ± 0.95 counts) (**Fig. 4h**). These findings are consistent with a deficit in the overall locomotor activity when the M_4_^+^ astrocytes are ablated.

### M_4_^+^ astrocytes do not regulate amphetamine hyperlocomotion

M_4_ mAChRs are known to counteract hyperlocomotion induced by amphetamine or the glutamate receptor agonist MK-801 (20, 24). This pivotal function in motor behaviours has been attributed to neuronal (presynaptic) M_4_ mAChR regulation of the dopamine release and signalling in the basal ganglia (25–28). Here, we examined if M_4_^+^ astrocytes might contribute to this by testing the efficacy of a M_4_ mAChR selective PAM, VU154, to reverse amphetamine-induced hyperlocomotion in Chrm4^CRE^:: Aldh1l1^DTA^ and littermate control mice. Administration of VU154 (10 mg/kg, i.p.) 20 mins before amphetamine (2 mg/kg, i.p.) significantly decreased amphetamine-induced hyperlocomotion in both groups of mice (**Fig. 4i, j**). Likewise, both genotypes displayed comparable measurements in mobility time **(Supplementary Fig. 4g)**. Together these data indicate that the role of M_4_^+^ astrocytes in locomotion observed earlier is independent from the dopamine-dependent motor function of M_4_ mAChR in the basal ganglia.

### M_4_ ^+^ astrocytes during prion disease

There is currently intense interest in the role of astrocytes in regulating the onset and progression of many neurodegenerative diseases and other brain disorders (29, 30). It is clear, however, that the responses to pathological stimuli of reactive astrocytes are very diverse, which could reflect the heterogeneous nature of astrocyte subpopulations with established differences in origin and temporal function (31). Here we probed the potential involvement of M_4_^+^ astrocytes in the pathology of murine prion disease, a progressive terminal neurodegenerative disease model associated with pronounced neuroinflammation and astrogliosis (32). Prion disease was induced in Chrm4^CRE^::MetRS* mice by inoculation with Rocky Mountain Laboratory (RML) prion-infected brain homogenate. At 20 weeks post-inoculation (w.p.i.), a sharp increase was observed in the number of M**_4_** ^+^ astrocytes (**Fig. 5a**), as determined by co-staining with the astrocytic marker GFAP **(Supplementary Fig. 5a)**. The lack of co-staining of these M_4_^+^ cells with microglial marker Iba1 excluded the possibility that these cells were infiltrating macrophages **(Supplementary Fig. 5b)**. In fact, the strong GFAP staining is consistent with these M_4_^+^ cells showing characteristics of reactive astrocytes, which was particularly evident in the region of the cerebellar nuclei (**Fig. 5b**).

**Fig 5.**
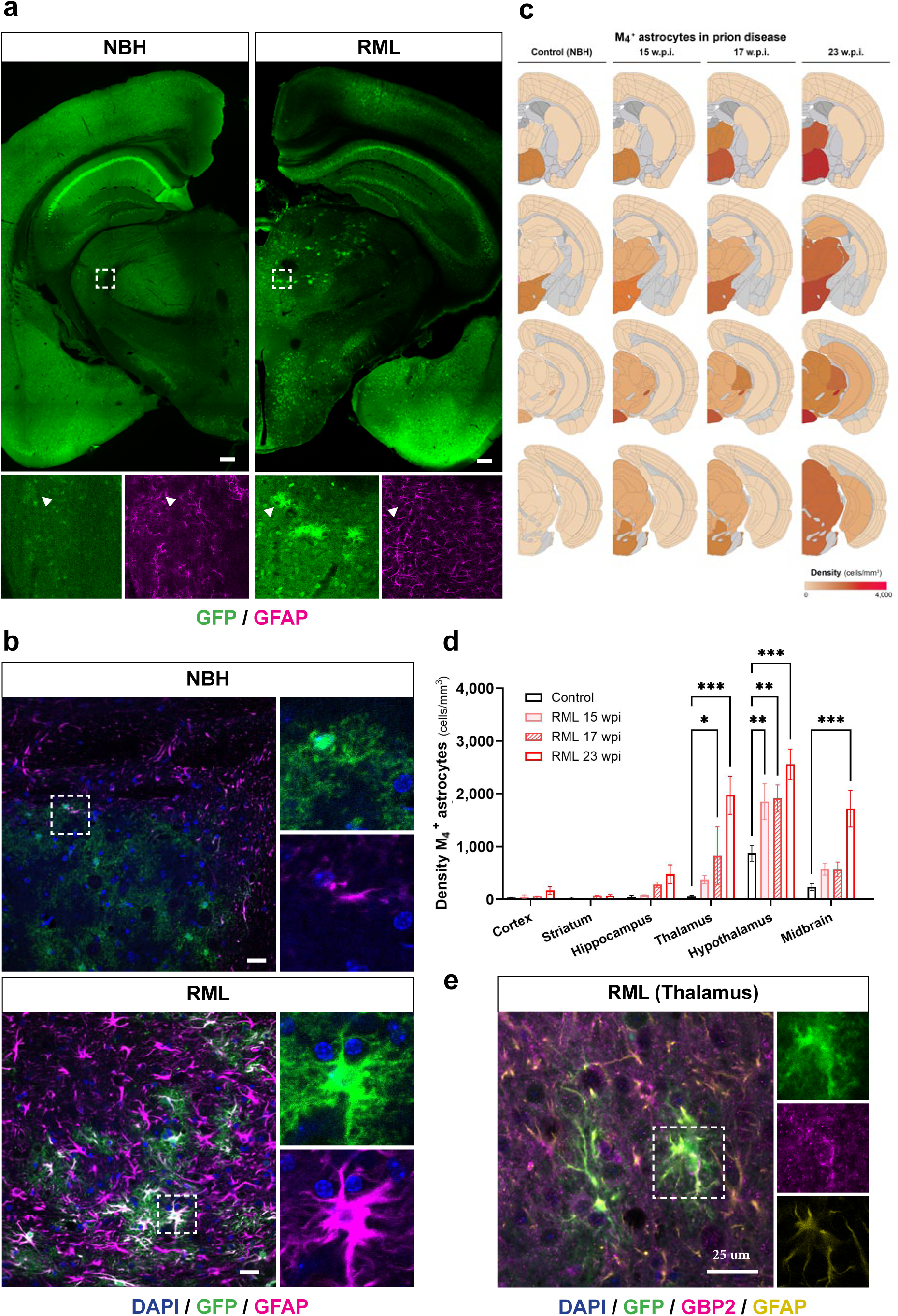
Regional and temporal expansion in the number of the M_4_^+^ astrocytes during mouse prion disease. **a**, Representative confocal images of GFP-labelled M_4_^+^ cells in coronal brain sections from Chrm4^CRE^::MetRS* animals after 20 w.p.i. of RML prion inoculation (right) or control animals treated with NBH (left). Panels below show a detail of M_4_^+^ astrocytes (arrowheads) co-stained with GFP (green) and GFAP (magenta) from the thalamic areas indicated with white square insets. **b**, Confocal images showing M_4_^+^ astrocytes co-stained with GFP (green) and GFAP (magenta) in the cerebellar nuclei of Chrm4^CRE^::MetRS* mice after 20 w.p.i. of RML prion disease (right) or NBH controls (left). **c**, Regional and temporal spreading pattern of the density of M_4_^+^ astrocytes at different stages of RML prion disease in Chrm4^CRE^::MetRS* mice plotted on brain anatomical maps. Darker red sections represent regions with higher densities in M_4_^+^ astrocytes, while not quantified areas are depicted in grey. **d**, Quantification of M_4_^+^ astrocytes density across brain regions in Chrm4^CRE^::MetRS* mice treated with RML prion after 15, 17 and 23 w.p.i. and compared to control animals inoculated with NBH (n = 2-3). **e**, Representative confocal images showing co-localisation of GFP (green), GFAP (magenta) and GBP2 in the hippocampus and thalamus of 20 w.p.i. RML-inoculated Chrm4^CRE^::MetRS* mice. Scale bar in **a** is 250 µm, and 25 µm in **b** and **e**. Data shown are represented as mean ± SEM. Statistical significance was calculated by two-way ANOVA followed by Bonferroni’s post hoc test (\*\*\**p* < 0.001, \*\**p* < 0.01).

A rostro-caudal examination of serial brain sections from RML-inoculated Chrm4^CRE^::MetRS* mice demonstrated that M_4_^+^ astrocytes were profoundly up-regulated in disease, even in regions such as the cortex, the hippocampus and the thalamus where these cells were not evident under normal “non-disease” conditions **(Supplementary Fig. 5a, b, c**). To interrogate regional differences, we estimated the number of M_4_^+^ astrocytes in each area during prion disease and displayed the cellular densities as heat maps overlaid onto the brain anatomy (**Fig. 5c)**. Our results indicated that the hypothalamus hosts the largest number of M_4_^+^ astrocytes. As prion disease is rapidly progressive, we further studied the evolution of these cellular subpopulations by comparing three time points after RML-inoculation (15, 18 and 23 w.p.i., **Fig. 5c, Supplementary Fig. 5d)**. We found that the number of M_4_^+^ reactive astrocytes gradually increased in almost all examined brain regions over time **(Fig. 5d)**. Interestingly, this prion-induced expansion was not observed in one of the biggest reservoirs of M_4_^+^ astrocytes, the cerebellar nuclei within the arbor vitae **(Supplementary Fig. 5e)**, although astrocytes in this region showed a prominent reactivity during disease **(Fig. 5b)**. Critically, M_4_^+^ astrocytes showed expression of GBP2 during prion disease **(Fig. 5e)**, a marker associated with a neurotoxic or pro-inflammatory phenotype (33). In addition to GBP2, these astrocytes also expressed metallothionein III (MT3) **(Supplementary Fig. 5f)**, which was similarly identified in our proteomic analysis as being upregulated upon activation of the M4 mAChR **(Supplementary Data 1)**. This spatio-temporal analysis of the M_4_^+^ astrocytes in prion disease indicated that these cells become activated in a pathological context and the number of reactive astrocytes that express the M_4_ mAChR continuously increases alongside the progression of the disease.

### Increased lifespan of prion disease inoculated mice where M_4_ ^+^ astrocytes are ablated

To establish the potential role played by M_4_^+^ astrocytes in prion disease, we evaluated the disease progression in mice where the M**_4_** ^+^ astrocytes had been ablated. Inoculation of both Chrm4^CRE^:: Aldh1l1^DTA^ and wild type littermate mice with RML resulted in a similar onset of prion disease and early/mid disease progression, as determined by near equivalent times when at least two early “clinical” signs of prion disease appeared **(Supplementary Fig. 6a)**. Similarly, there was equivalence in progressive weight loss **(Supplementary Fig. 6b)**, decreased motor co-ordination and decline in burrowing behaviour (a sign of hippocampal function) between the genotypes **(Fig. 6a, b)**. Importantly, despite a similar progression of early and mid-stages of disease, Chrm4^CRE^:: Aldh1l1^DTA^ mice showed extended lifespan compared to wild type littermates (mean survival 164.7 ± 0.9 and 158.6 ± 1.5 days, respectively, \*\**p* < 0.01) **(Fig. 6c).** This survival effect was more prominent in females **(Fig. 6d)**, which could be explained by their shorter incubation periods to Prp^Sc^ compared to males **(Fig. 6e)** when exposed to the same quantity of infectious material (34). Our result emphasises the importance of sex as a biological variable in prion pathogenesis, previously observed in mice depleted of NG2 glia (35) and in human sporadic Creutzfeldt-Jakob disease (sCJD) (36).

**Fig. 6.**
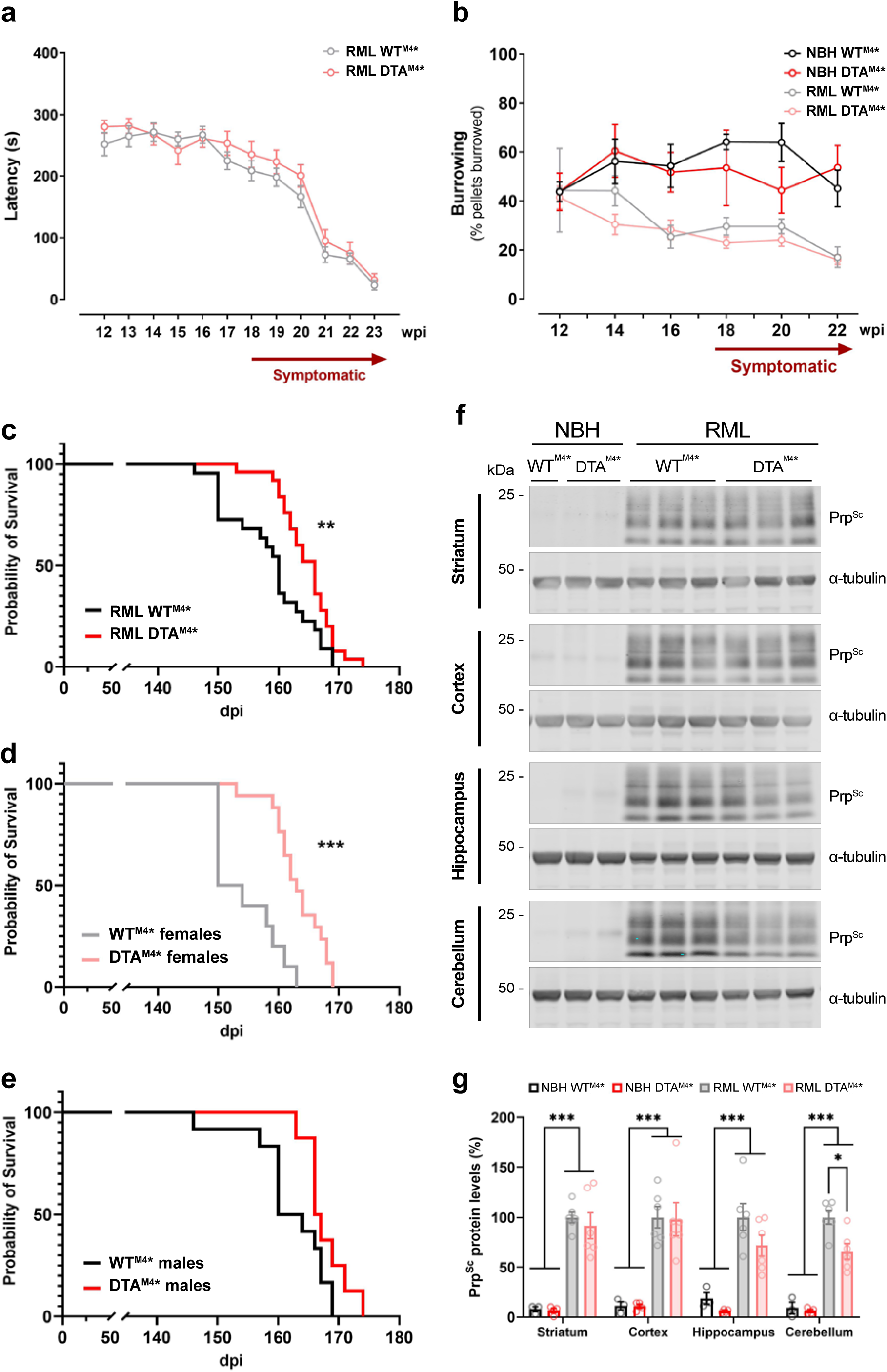
Ablation of M_4_^+^ astrocytes prolongs survival and reduces Prp^Sc^ deposition in the cerebellum of prion-infected mice. **a**, Evaluation of progressive motor deficits by rotarod in prion-diseased DTA^M4^* and WT^M4^* mice from 12 w.p.i (n = 8-12). **b**, Burrowing behaviour (percentage of food pellets displaced from the tube) of control or prion-infected DTA^M4^* and WT^M4^* mice assessed from 12 w.p.i. (n = 5 for NBH, n = 10-12 for RML). **c**, Kaplan-Meier plot showing a significant increase in survival probability for RML-prion inoculated mice after ablation of M_4_^+^ astrocytes (DTA^M4^*) compared to wild-type littermates (WT^M4^*) (n = 22 - 25). Sex-differences are observed when survival curves are compared between females (**d**, n = 10-17) and males (**e**, n = 8-12). Curves were analysed with a Gehan-Breslow-Wilcoxon test (\*\**p* < 0.01, \*\*\**p* < 0.001). **f**, Immunoblot analysis of PK-resistant Prp^Sc^ from the striatum, cortex, hippocampus and cerebellum of DTA^M4^* and WT^M4^* mice inoculated with NBH or RML-prion. Evaluated samples were collected at terminal stages of prion diseased mice with equivalent survival times. **g**, Quantification of Prp^Sc^ shows accumulation of the PK-resistant protein in all evaluated brain regions from diseased animals. Ablation of M_4_^+^ astrocytes significantly reduces the deposition of Prp^Sc^ in the cerebellum. Graph represents means ± SEM of Prp^Sc^ levels relative to α-tubulin expression (n = 4-6) expressed as a percentage of the values obtained for RML-prion diseased wild-type mice (WT^M4^*). Statistical significance was calculated by two-way ANOVA followed by Tukey’s post-hoc test (\**p* < 0.05, \*\*\**p* < 0.001).

We next assessed the accumulation of misfolded, insoluble scrapie prion protein (Prp^Sc^), which constitutes the major component of infectious prion disease. At the terminal stage, a dramatic deposition of Prp^Sc^ was detected in the hippocampus, cortex, striatum and cerebellum of prion-inoculated mice when compared to the non-diseased controls **(Fig. 6f, g)**. Importantly, the accumulation of Prp^Sc^ was significantly decreased in prion inoculated Chrm4^CRE^:: Aldh1l1^DTA^ compared to wild type controls in the cerebellum, but not in the rest of the brain regions **(Fig. 6g)**. However, this reduction was not accompanied by changes in other indicators of neurodegeneration (ApoE and serpinA3N) or markers of neuroinflammation (GFAP, Iba1 and clusterin) **(Supplementary Fig. 6c, d)**. Overall, these data indicate that the ablation of M_4_^+^ astrocytes does not affect the onset of the clinical symptoms or progression of prion disease but extends the lifespan of mice, indicating a potential detrimental effect of M_4_^+^ astrocytes in the later stages of disease.

## Discussion

Despite the considerable progress in understanding the neuronal populations that display the M_4_ mAChR (37–39), the identity, location and function of non-neuronal CNS cells, in particular glial cells, expressing this receptor subtype have remained largely unexplored. Here, we reveal a discrete subpopulation of M_4_ mAChR expressing astrocytes dispersed across several areas of the mouse CNS. Genetic ablation of these astrocytes demonstrated a role in neurological responses associated with locomotion, while other M_4_ mAChR-associated processes such as anxiety and the regulation of responses to amphetamine remained unaffected. Importantly, there was a dramatic expansion of this subpopulation of astrocytes during the progression of neurodegeneration in a model of prion disease. Ablation studies indicated a detrimental role for these M**_4_** ^+^ astrocytes in later disease stages. In these ways our study contributes to a growing awareness that astrocytes are not uniform but rather a diverse set of cell types with varying cellular and functional characteristics. Appreciating the nature and complexity of the various subtypes of astrocytes will be essential if we are to understand their role in normal neurophysiology and if we are to target these cells in brain diseases.

Under physiological conditions, we found M**_4_** ^+^ astrocytes in the hypothalamus and dorsal parts of the brainstem. These regions are critical regulators of energy balance and appetite, and astrocytes in these areas have been reported to contribute to these tasks by modulating the activity of adjacent neurons (40). Thus, manipulation of hypothalamic and brainstem astrocytes has been reported to influence metabolic processes including feeding and glucose metabolism (41–46), as well as circadian rhythm, and sleep (47, 48). Interestingly, ablation of M_4_^+^ astrocytes in mice did not result in alterations in the weight, body temperature or daily activity cycle. It was, however, of relevance that M_4_^+^ astrocytes regulate locomotion, an activity consistent with the expression of the receptor in a diverse array of the cerebellar astrocytes, which are activated during voluntary and enforced locomotion (49, 50). It is important to note that M_4_^+^ astrocytes are involved in specific aspects of motor control whilst other behaviours such as balance and coordination, typically altered after a loss of Bergmann glia support (23), remained normal in M_4_^+^ astrocyte-depleted mice. Furthermore, the ablation of M_4_^+^ astrocytes had no impact on basal ganglia-mediated hyperlocomotion responses, classically linked with neuronal M_4_ mAChRs (25–28), despite the fact that astrocytes in this region can modulate these psychomotor behavioural effects (51). Hence, our studies have dissected a locomotion response that may be under the regulation of M_4_ mAChR expressed on a subpopulation of astrocytes.

While our study primarily identified the existence of a novel population of astrocytes, the neuro-anatomical distribution of these cells, and the function that these astrocytes might play under normal and neurodegenerative disease scenarios, it is a particular challenge to relate this to the activity of M_4_ mAChRs themselves. We started to investigate this in primary cultures where we used a genetic approach that allows for the signalling and proteomic interrogation of the M_4_^+^ cerebellar astrocytes amongst a mixed population of astrocytes. This has provided an in-depth proteomic profile of M_4_^+^ astrocyte subpopulation and demonstrated that selective M_4_ mAChR activation resulted in the regulation of key cytoskeletal, metabolic and translational proteins that can significantly impact astrocyte biology and activation status. For example, profilin-2 can modulate astrocytic morphology (52); MT3 plays a role in the clearance of the prion-like amyloid β (53), and the low-density lipoprotein receptor-related protein-1 (LRP1) is important for the astrocyte-neuron communication (54, 55). Interestingly, LRP1 is downregulated together with M_4_ mAChR in the cerebellum of patients with Rett syndrome (56), and mouse models show motor deficits when the *MeCP2* gene is deleted from the cerebellum but not when is restricted to Purkinje or granular neurons (57), highlighting the contribution of the cerebellar astrocyte pool.

The involvement of astrocytes, and indeed microglia, in protection against neurodegenerative diseases is an extremely active area of research (58). Astrogliosis is a widespread response in prion disease, although each brain region contains reactive astrocytes with heterogenous phenotypes and functions determined by the microenvironment (59). Our study showed a striking increase in M_4_^+^ astrocytes in the thalamus and hypothalamus, brain regions identified as particularly susceptible to prion deposition and neuroinflammatory cascades in RML and other prion strain models (60, 61). Astrocytes are known to adopt either neuroprotective or neurotoxic phenotypes in response to pathological stimuli. While their early activation may support neuronal survival and clearance of misfolded proteins, functions that are protective in diseases like Alzheimer’s disease (62), the prolonged activation of astrocytes is widely considered to be detrimental (63, 64). This certainly seems to be the case for M_4_^+^ astrocytes which do not seem to make a significant impact on early-and mid-stages of disease but rather are involved in acceleration of the later stages of disease. The expression of GBP2 in M_4_⁺ astrocytes, a marker linked to neurotoxic reactivity, further supports their detrimental role in late-stage disease. How the M_4_ mAChR itself is involved in the regulation of these astrocytes at latter disease stages, and whether agonists or antagonists might make an impact is at present uncertain. What is clear, however, is that our findings support the selective targeting of neurotoxic astrocytes while preserving glial subtypes with neuroprotective roles as a promising therapeutic strategy for prion disease. A therapeutic option previously discarded after observing that preventing the formation of the neurotoxic C3⁺ astrocytes in mice accelerated disease progression (33). This dichotomy highlights the delicate balance between protective and harmful astrocyte states and the need for careful and thorough characterisation of these astrocytic subpopulations.

In conclusion by identifying a previously unappreciated astrocyte subpopulation we not only provide further evidence of the diversity of astrocytes but by ablation of M_4_^+^ astrocytes highlight the role of this astrocyte population in the regulation of locomotion and neurodegenerative disease pathology.

## Materials and Methods

### Animals

Mice were maintained on a 12-h light–dark cycle with ad libitum standard mouse chow food and water.

### Generation of Chrm4^CRE^ mice

Transgenic Chrm4^CRE^ mice were generated by genOway (Lyon, France). Briefly, a CRE-IRES cassette was inserted in frame of the ATG in exon 2 of the endogenous *Chrm4* gene, by homologous recombination in embryonic stem (ES) cells. To this end a targeting vector was created containing long and short homology arms to the *Chrm4* genomic region. The targeted vector was completed with an FRT-flanked neomycin resistance gene upstream the CRE sequence to select ES cells that had undergone homologous recombination, and a diphtheria toxin cassette upstream the long homology arm for negative selection. The linearised targeting construct was transfected into C57BL/6N ES cells by electroporation. Positive clones were assessed by PCR and whole locus sequence analysis to validate the presence of the correct recombination event before being injected into blastocysts derived from C57BL/6J female mice. The resulting highly chimeric males (chimerism rate above 50%) were then crossed to a FLP recombinase-expressing deleter mouse strain to generate heterozygous knock-in animals devoid of neomycin. Genotyping was initially confirmed by PCR on genomic DNA obtained from ear biopsies using the primers 5’ CACTGCCCACCTGTCTCCAG and 5’ GGTTGTGGGCTGTTGTGACCAG, which flank the insertion site of the CRE-IRES cassette; and further confirmed with the pair of primers 5’ GCTAGAGCCTGTTTTGCACG and 5’ GCTAGAGCCTGTTTTGCACG, which amplify a region of the CRE recombinase sequence. PCR-positive animals were further validated by whole locus sequencing. The results were later corroborated by fluorescent quantitative PCR (qPCR) using a commercial genotyping service (Transnetyx, Cordova, TN). This service also analysed further samples to verify the genotype of the first litter of every new breeder pair for the maintenance of the Chrm4^CRE^ colony.

Homozygous Chrm4^CRE^ were crossed with homozygous R26-CAG-GFP,-MetRS* (Jackson Laboratory, strain: 028071) to generate Chrm4^CRE^::MetRS*, and heterozygosity for both alleles was confirmed by Transnetyx genotyping service. Chrm4^CRE^ animals were mated with hemizygous Aldh1l1-eGFP,-DTA mice (Jackson Laboratory, strain: 026033) to produce Chrm4^CRE^::Aldh1l1^WT^ and Chrm4^CRE^:: Aldh1l1^DTA^. All these animals were genotyped with specific probes designed for CRE recombinase, eGFP and Chrm4 sequences at weaning by a commercial vendor (Transnetyx). Chrm4^CRE^:: Aldh1l1^DTA^ and wild type littermate animals were kept mixed in housing cages, except when measuring the food and water intake and during telemetry monitoring experiments. The generation of M_4_ KO mice has been described elsewhere (65).

### Measurement of organ weight, food and water consumption

Chrm4^CRE^:: Aldh1l1^DTA^ and wild type littermate mice were housed separated by genotype and food and water intakes were measured every 3 days for 1-2 weeks. Average daily food and water consumption per animal were calculated and no differences were observed between genotypes or sex.

The brain, liver, heart, spleen and kidneys from Chrm4^CRE^:: Aldh1l1^DTA^ and wild type littermate animals were collected (n=3-5, 4-5 months old) and the weight was recorded and adjusted for body weight. Organ dimensions were also measured, and no differences were observed (data not shown).

### Rotarod

Motor performance was tested using the 5-lane rotarod (Cat. #47650, Ugo Basile, Germany). Mice were weighed prior every testing, which happened weekly for both prion infected and non-infected 10-week-old male mice for 15-20 weeks. Each day, mice were placed on the 3 cm diameter rod for 2 habituation phases, consisting of a constant rotation speed at 4 rpm for 90 s. For the testing phase, the animals were placed on the rod that accelerated from 4 rpm to 40 rpm over a period of 5 min. The mice were tested 2 times with a rest period of at least 15 min between each trial. Animals that fell before the rod reached 15 rpm speed were placed back on the beam, but if they fell more than once in that period, they were returned to the home cage. If mice turned on the rod, the test was stopped, and the time recorded. The latency to fall (time that the mouse remains on the rod) was registered automatically by a trip switch under the floor of each rotating drum, and the result was used as an indicator of motor coordination. The activity of Chrm4^CRE^:: Aldh1l1^WT^ in this test was comparable to wild type animals with a C57BL/6 background (data not shown).

### Elevated Plus Maze (EPM)

The EPM test was used to assess anxiety-related behaviour in mice. 9-10 weeks old male and female animals were placed in the centre of a cross shaped maze (arms of 50 x 10 cm), with 2 enclosed arms (black walls of 30 cm height) and 2 dimly illuminated open arms opposite each other. The test was performed before the start of the light cycle and lasted for 5 min. The time spent in the open arms versus the enclosed spaces versus, the immobility time and the number of entries to each arm were recorded using the ANY-maze video tracking software (Stoelting).

### Open field

The open field activity of 8-12 weeks old male and female mice was monitored using a clear Perspex square box (50 × 50 cm). Animals were placed in the centre of the illuminated open field arena and allowed to freely explore for 10 minutes. The time spent in the centre of the arena, the distance travelled and speed were assessed using the ANY-maze tracking system (Stoelting).

### Hindlimb clasping test

The degree of hindlimb clasping as an indicator of motor dysfunction was evaluated in 8-9 weeks old mice. Animals were suspended by their tails for 10-20 s and rated from 0 to 3 according to the following score system: 01 corresponded to hindlimbs consistently splayed outward and away from the abdomen; 11 was assigned when 1one hindlimb was retracted inwards towards the abdomen for more than 50% of the time; a score of 2 1meant that both hindlimbs were partially retracted inwards towards the abdomen for at least 50% the time; and 311was given when both hindlimbs were entirely retracted inwards towards the abdomen for at least 50% of the observation period (66). Two separate trials were performed per animal. All evaluated Chrm4^CRE^:: Aldh1l1^DTA^ and Chrm4^CRE^:: Aldh1l1^WT^ animals showed a consistent score of 0 in this test.

### Ledge test

Mice aged 8-9 weeks old were placed on the cage ledge and recorded to measure balance and coordination. Paw placement and forward movement as it walked along the ledge were observed and scored from 0 to 3 as in (66): 0, for normal walking; 1, if the mouse lost its footing; 2, if the hindlimbs were not used, and 3, if the animal fell off the ledge. Data from three trials were averaged for each mouse and none of the Chrm4^CRE^:: Aldh1l1^DTA^ and Chrm4^CRE^:: Aldh1l1^WT^ mice presented any deficits in balance and coordination.

### Telemetry

A TA-F10 temperature and activity probe (Data Sciences International) was implanted in the abdominal cavity of 9-13 weeks old male and female Chrm4^CRE^:: Aldh1l1^DTA^ and wild type littermate mice (n=8, 19-30 g), using aseptic techniques under a surgical level of isoflurane anaesthesia. Carprofen (10 mg/kg) and buprenorphine (0.1 mg/kg) were s.c. administered 30 min before surgery for pain relief. After surgery, mice were housed individually, and post-operative analgesia was given 6 h (buprenorphine) and 24 h later (carprofen and buprenorphine). One week after surgery, when the animals completely recovered, the cages were placed over the receiver panels and the TA-F10 transmitters were activated. Activity and body temperature data were sampled every 1 min for 10 s and continuously acquired during 4 consecutive days using the Ponemah software (Data Sciences International). The measurements from the final 3 days were averaged together for each mouse. No differences were observed between males and females.

### Amphetamine-induced hyperlocomotion

Male and female Chrm4^CRE^:: Aldh1l1^DTA^ and wild type littermate mice (n=6-7, 10-11 weeks old) were placed in an open field arena and the animals were tracked with the ANY-maze software (Stoelting). Habituation lasted for 20 min before being i.p. injected with vehicle (10% Tween 80 in sterile water) or VU0467154 (10 mg/kg). A dose of D-amphetamine (2 mg/kg in saline, i.p.) was administered 20 min later, and locomotor activity was recorded for an additional 90 min. The mobility and distance travelled over time were expressed and averaged per 5 min bins. Total activity data were calculated as the total distance travelled 60 min after the time of D-amphetamine administration.

### Burrowing

The voluntary behaviour of removal material from an enclosed tube was assessed in control and prion-infected Chrm4^CRE^:: Aldh1l1^DTA^ and wild type littermate female mice every week from 9 w.p.i. until survival time. Mice were placed into individual cages (22 × 36 cm) with an acrylic clear tube filled with food pellets for 2 h. The burrowing activity was calculated by subtracting the weight of the remaining pellets at the end of the experiment from the starting weight (140 g). Mice were acclimatised to the burrowing cage the week before the start of the test.

### Prion mouse model

Male and female Chrm4^CRE^::MetRS*, Chrm4^CRE^:: Aldh1l1^DTA^ and Chrm4^CRE^:: Aldh1l1^WT^ mice aged 3-4 weeks old were i.c. inoculated, under a surgical level of isoflurane anaesthesia, into the right brain parietal lobe with 30 µl of either 1% NBH (Normal Brain Homogenate) or 1% RML prion (Rocky Mountain Laboratories), as previously described (67, 68).

### Evaluation of prion disease progression and survival

Chrm4^CRE^:: Aldh1l1^DTA^ and Chrm4^CRE^:: Aldh1l1^WT^ control and prion-infected mice were co-housed in mixed-genotype cages to minimise environmental and housing-related bias. Clinical assessment of prion disease progression was performed daily by a trained technician who was blinded to the genotype of the animals, and the scoring protocol followed established criteria consistent with previously published methodologies (7, 68, 69). Disease progression was evaluated according to the presence of early indicator and confirmatory signs of prion disease. Early indicator signs included piloerection, mild loss of coordination, erect penis, clasping of hindlimbs when lifted by tail, intermittent generalised tremors, sustained erect ears, rigid tail, discontinued hunched posture, or subdued behaviour. Confirmatory signs of prion disease included righting reflex impairment, sustained hunched posture, dragging of limbs, and/or significant abnormal breathing. Animals were humanely killed when they showed two confirmatory signs or two early indicators plus one confirmatory sign, and the days after inoculation were considered as the survival time.

### Preparation of protein extracts and immunoblot analysis

Mice were culled via cervical dislocation, and different regions of the brain were dissected and snap frozen on dry ice. Samples were lysed in RIPA buffer (10 mM Tris-Cl, 1 mM EDTA, 1% Triton X-100, 0.1% sodium deoxycholate, 0.1% SDS, 140 mM NaCl, pH 7.5) supplemented with protease inhibitor tablets (Cat. #11697498001, Roche) and incubated for 30 min at 4 °C. Lysates were then briefly sonicated (Cat. #12337338, Fisherbrand) with 2 pulses of 15 s at an amplitude of 40% (∼50 W) until fully dissolved. Samples were then centrifuged at 10,000 × g for 10 min, supernatants collected, and protein concentration calculated using a BCA assay kit (Cat. #23225, Pierce). Equal concentrations of protein samples were resuspended in Laemmli loading buffer (10% SDS, 300 mM Tris-HCl pH 7.2, 0.05% bromophenol blue, 10% β-mercaptoethanol) and heated at 60°C for 10 min. For the assessment of PrP^Sc^ levels, 60 μg of each lysate were treated with proteinase K (10 μg/ml, Cat. P2308, Sigma-Aldrich) for 10 min at 37°C. The reaction was stopped by the addition of Laemmli sample buffer and boiling at 95°C for 5 min. 20-40 μg of each cell lysate were resolved on NuPAGE precast 4-12% gels (Cat. #NP0321, Invitrogen) followed by transfer onto nitrocellulose membranes (Cat. #88018, Thermo Scientific). Protein transfer was verified with Ponceau-S staining (0.2% Ponceau-S red, 1% acetic acid), before membranes were blocked with 5% non-fat dry milk in TBST (25 mM Tris-HCl; pH 7.6, 100 mM NaCl, 0.5% Tween 20) for 1 h at room temperature. Membranes were incubated with the corresponding primary antibodies in the blocking solution overnight at 4°C. After washing three times for 5 min each with TBST, secondary antibodies were added to the membranes for 1 h at RT. Antibodies used in this study are detailed in Tables 1 and 2. Immunoreactive bands were visualised using the Licor Odyssey system and densitometry of fluorescence was measured using Image Studio Lite (Licor Biosciences).

**Table 1:**
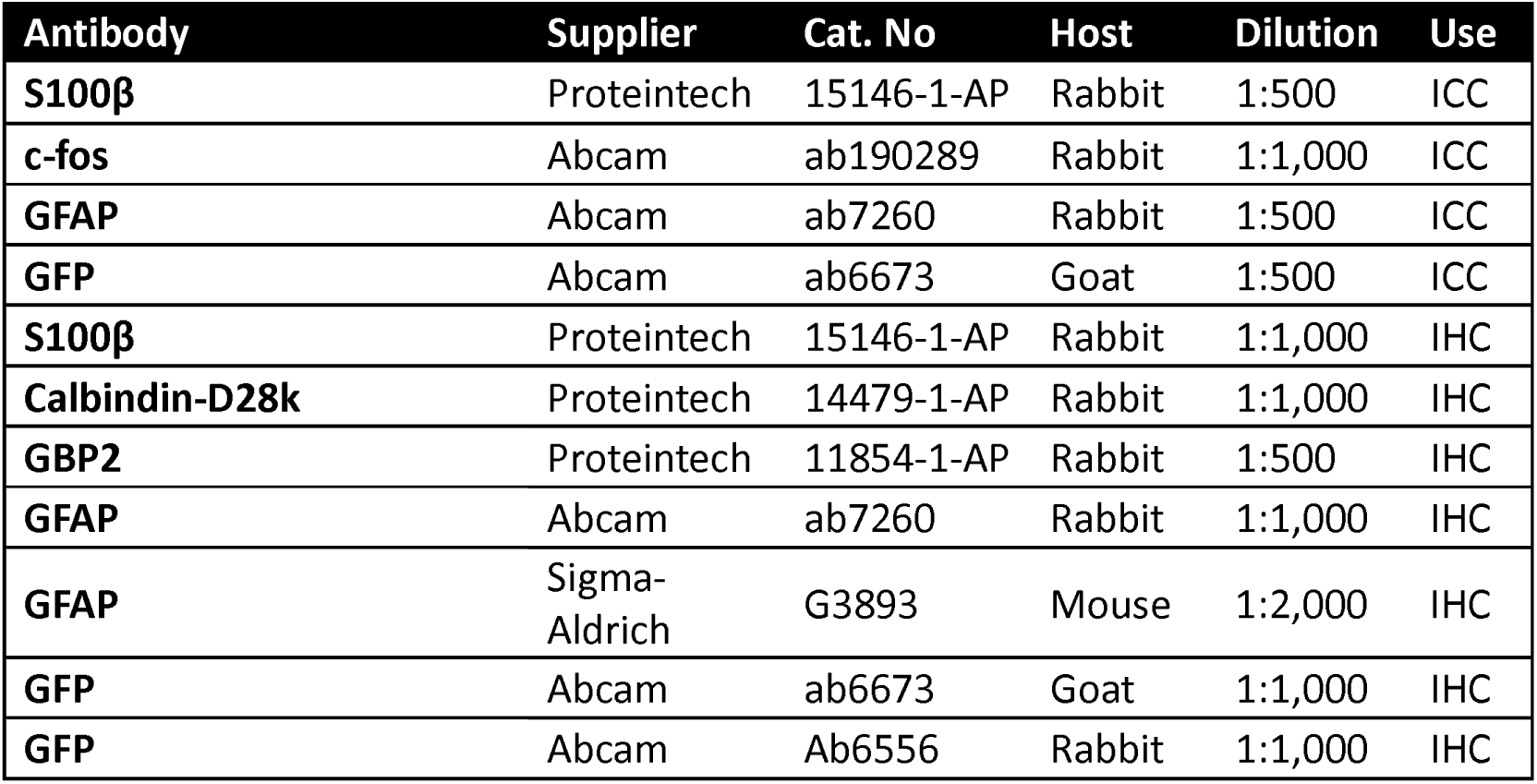
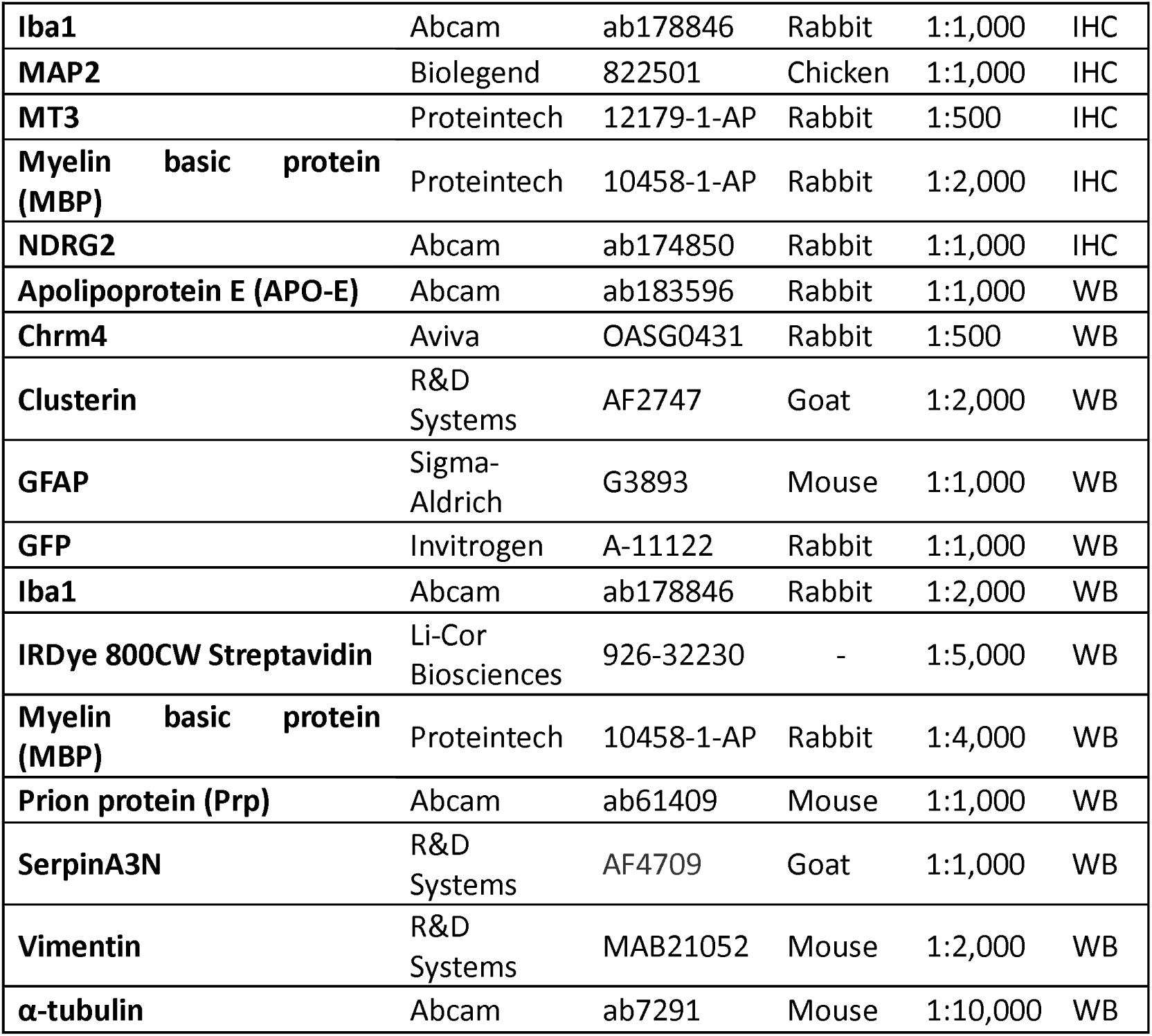
Primary antibodies. ICC, immunocytochemistry; IHC, immunohistochemistry; WB, western blot.

| Antibody | Supplier | Cat. No | Host | Dilution | Use |
| --- | --- | --- | --- | --- | --- |
| **S100β** | Proteintech | 15146-1-AP | Rabbit | 1:500 | ICC |
| **c-fos** | Abcam | ab190289 | Rabbit | 1:1,000 | ICC |
| **GFAP** | Abcam | ab7260 | Rabbit | 1:500 | ICC |
| **GFP** | Abcam | ab6673 | Goat | 1:500 | ICC |
| **S100β** | Proteintech | 15146-1-AP | Rabbit | 1:1,000 | IHC |
| **Calbindin-D28k** | Proteintech | 14479-1-AP | Rabbit | 1:1,000 | IHC |
| **GBP2** | Proteintech | 11854-1-AP | Rabbit | 1:500 | IHC |
| **GFAP** | Abcam | ab7260 | Rabbit | 1:1,000 | IHC |
| **GFAP** | Sigma-Aldrich | G3893 | Mouse | 1:2,000 | IHC |
| **GFP** | Abcam | ab6673 | Goat | 1:1,000 | IHC |
| **GFP** | Abcam | Ab6556 | Rabbit | 1:1,000 | IHC |
| **Iba1** | Abcam | ab178846 | Rabbit | 1:1,000 | IHC |
| **MAP2** | Biologend | 822501 | Chicken | 1:1,000 | IHC |
| **MT3** | Proteintech | 12179-1-AP | Rabbit | 1:500 | IHC |
| **Myelin basic protein (MBP)** | Proteintech | 10458-1-AP | Rabbit | 1:2,000 | IHC |
| **NDRG2** | Abcam | ab174850 | Rabbit | 1:1,000 | IHC |
| **Apolipoprotein E (APO-E)** | Abcam | ab183596 | Rabbit | 1:1,000 | WB |
| **Chrm4** | Aviva | OASG0431 | Rabbit | 1:500 | WB |
| **Clusterin** | R&D Systems | AF2747 | Goat | 1:2,000 | WB |
| **GFAP** | Sigma-Aldrich | G3893 | Mouse | 1:1,000 | WB |
| **GFP** | Invitrogen | A-11122 | Rabbit | 1:1,000 | WB |
| **Iba1** | Abcam | ab178846 | Rabbit | 1:2,000 | WB |
| **IRDye 800CW Streptavidin** | Li-Cor Biosciences | 926-32230 | - | 1:5,000 | WB |
| **Myelin basic protein (MBP)** | Proteintech | 10458-1-AP | Rabbit | 1:4,000 | WB |
| **Prion protein (Prp)** | Abcam | ab61409 | Mouse | 1:1,000 | WB |
| **SerpinA3N** | R&D Systems | AF4709 | Goat | 1:1,000 | WB |
| **Vimentin** | R&D Systems | MAB21052 | Mouse | 1:2,000 | WB |
| **α-tubulin** | Abcam | ab7291 | Mouse | 1:10,000 | WB |

**Table 2:** Secondary antibodies. ICC, immunocytochemistry; IHC, immunohistochemistry; WB, western blot.

| **Antibody** | **Supplier** | **Cat. No** | **Host** | **Dilution** | **Use** |
| --- | --- | --- | --- | --- | --- |
| **Anti-Rabbit Alexa Fluor Plus 647** | Invitrogen | A32795 | Donkey | 1:500 | ICC |
| **Anti-Chicken Alexa Fluor 594** | Invitrogen | A11042 | Goat | 1:1,000 | IHC |
| **Anti-Rabbit Alexa Fluor 488** | Invitrogen | A32731 | Goat | 1:1,000 | IHC |
| **Anti-Goat Alexa Fluor Plus 488** | Invitrogen | A32814 | Donkey | 1:1,000/<br>1:500 | IHC/<br>ICC |
| **Anti-Mouse Alexa Fluor Plus 594** | Invitrogen | A32744 | Donkey | 1:1,000/<br>1:500 | IHC/<br>ICC |
| **Anti-Rabbit Alexa Fluor Plus 594** | Invitrogen | A32754 | Donkey | 1:1,000/<br>1:500 | IHC/<br>ICC |
| **IRDye 680RD Anti-Goat** | Li-Cor Biosciences | 925-68074 | Donkey | 1:5,000 | WB |
| **IRDye 680RD Anti-Mouse** | Li-Cor Biosciences | 925-68072 | Donkey | 1:5,000 | WB/<br>IHC |
| **IRDye 800CW Anti-Mouse** | Li-Cor Biosciences | 926-32212 | Donkey | 1:5,000 | WB |
| **IRDye 800CW Anti-Rabbit** | Li-Cor Biosciences | 925-32213 | Donkey | 1:5,000 | WB |

### Astrocyte cultures

Murine astrocyte cultures were performed according to the AWESAM protocol (70) with some modifications. Brains from Chrm4^CRE^::MetRS* or WT pups (P0-P5) were collected and immediately placed in ice cold HBSS (ThermoFisher Scientific). After removing of meninges, the cerebellum or the cerebrum were dissected and incubated with 0.05% trypsin-EDTA (Cat. #25300054, ThermoFisher Scientific) at 37°C for 10 min followed by gentle mechanical dissociation. Cells were centrifuged at 1,000 x g for 3 min, resuspended in high glucose, Glutamax-supplemented DMEM (Cat. #61965026, Gibco) with 10% Foetal Bovine Serum (Cat. #10500064, Gibco), 100 units/ml penicillin and 100 µg/ml streptomycin (Cat. # 15070063, Gibco), and plated on flasks precoated with poly-D-lysine (100 µg/ml, Cat. #P0899, Sigma-Aldrich). After 7 days in vitro (DIV), the flasks were shaken at 180 rpm for 30 min at 37°C, media changed and further shaken at 240 rpm for 6 h. Astrocytes were detached and seeded at a density of 0.2 x 10^6^ cells/ml on the corresponding experimental poly-D-lysine plates. After a couple of days, media was replaced with NB+H media, containing Neurobasal (Cat. # 21103049, Gibco) supplemented with B-27 (Cat. #17504044, Gibco), 2 mM Glutamax (Cat. # 35050061, Gibco), 10 ng/mL heparin-binding EGF-like growth factor (Cat. #E4643, Sigma-Aldrich), 100 units/ml penicillin and 100 µg/ml streptomycin for 5 days before the start of the experiments.

### ERK_1/2_ phosphorylation assay

Cultured primary astrocytes from WT mice were seeded onto 96 well plates precoated with poly-D-lysine at a density of 20,000 cells/well in NB+H media. One week later, the media was replaced, and cells were incubated with 80 µl of Neurobasal without additives for 4 h. Cells were treated for 10 min with vehicle or 10 µM of the M_4_ mAChR antagonist VU6028418 before stimulation with the indicated concentrations of oxotremorine for 5 min at 37°C in a final volume of 100 µl. Maximum ERK_1/2_ phosphorylation response was obtained by treating the cells with FBS. ERK phosphorylation was evaluated using a Homogenous Time-Resolved FRET (HTRF)-based phospho-ERK (Thr202/Tyr204) cellular kit (Cat. #64ERKPEH, Revvity). Briefly, lysis buffer was added immediately after the end of the stimulation and incubated for 30 min. After homogenisation, 16 μl of the lysate were transferred to a 384-well white ProxiPlate (PerkinElmer) and combined with 4 μl HTRF premixed antibodies. for 4 h at room temperature. Fluorescence emission at 665 and 620 nm was measured after 4 h with a PHERAstar plate reader (BMG Labtech) and results were normalised to the response stimulated by FBS.

### Immunocytochemistry analysis

Cerebral or cerebellar primary astrocytes from Chrm4^CRE^::MetRS* mice were grown onto 24 well plates with 13 mm glass coverslips previously treated with poly-D-lysine as indicated above. Cells were fixed with 4% formaldehyde for 1 h at room temperature. Coverslips were then washed 3 times with PBS and blocked with PBS supplemented with 1% BSA and 0.2% Triton X-100 for 1 h. Subsequently, astrocytes were incubated with the indicated primary antibodies diluted in blocking solution overnight at 4°C in a humidity chamber. Detection was achieved using appropriate secondary antibodies conjugated to Alexa Fluor 488 or Alexa Fluor 594 (1:500, Life Technologies) and coverslips were mounted on glass slides using Fluoromount-G with DAPI (ThermoFisher Scientific). Confocal images were acquired with a Zeiss Pascal laser-scanning confocal microscope (LSM) 880 using the LSM image acquisition software.

For analysis of c-fos expression, the NB+H media was replaced with non-supplemented Neurobasal, and cells were incubated at 37°C and 5% CO_2_ for 4 h, before being stimulated with vehicle or 10 µM oxotremorine for 60 min. To evaluate M_4_ mAChR action, cultures were simultaneously treated with vehicle, 10 µM VU0467154 or 5 µM VU6028418. Immunocytochemistry was performed as described above and, for each replicate (n = 3), 8 confocal images (425 µm x 425 µm) per condition were capture, randomly selected using DAPI staining from areas with equivalent number of cells. The images were analysed using ImageJ Fiji (NIH Image). Automatic quantification of c-fos^+^ cells was performed using the “Analyze particle” plugin. M_4_ mAChR-expressing astrocytes were manually counted and M_4_^+^/c-fos^+^ double stained cells were noted. Total number of cells were also counted automatically with the software tools. The experimental conditions were blinded to the investigator performing the image analysis. This methodology was also used to calculate the proportion of M_4_^+^ astrocytes present in cerebral and cerebellar cultures.

For FUNCAT analysis, Chrm4^CRE^::MetRS* astrocytes were incubated with 4 mM L-Azidonorleucine hydrochloride (ANL, Tocris) for 24 h. Cells were fixed and blocked as previously described, followed by click chemistry to visualize the ANL-labelled newly synthesized proteins. The Cu(I)-catalysed azide/alkyne cycloaddition (CuAAC) reaction comprised the addition of 0.45 mM CuSO_4_, 0.9 mM Tris((1-hydroxy-propyl-1H-1,2,3-triazol-4-yl) methyl) amine (THPTA, Cat. #CLK-1010, Jena Bioscience), 0.2 mM Cy3-alkyne (Cat. #777331, Sigma-Aldrich) and 5 mM Sodium ascorbate to the cells for 24 h. Coverslips were washed with PBS containing 0.5 mM EDTA for 10 min, followed by 3 washes with blocking solution and the remaining steps of the immunocytochemistry proceeded as indicated above.

### Immunohistochemistry

Mice were deeply anaesthetised with isofluorane and then they were transcardially perfused with 20 ml saline solution followed by 20 ml 4% formaldehyde in PBS. Brains were removed and incubated overnight in the same fixative. The whole brain was coronally or sagittally sectioned (50 μm) using a VT1200S vibratome (Leica). After blocking the free-floating sections with PBS supplemented with 0.5% Triton X-100 and 1% bovine serum albumin for 1 h at room temperature, they were then incubated with the indicated primary antibodies diluted in blocking solution 24 to 48 h at 4°C. Antibodies used in this study are listed in Tables 1 and 2. Brain sections were washed 3 times with PBS, followed by incubation with the corresponding secondary antibodies diluted in blocking solution for 4 h at room temperature. Sections were placed onto 2% gelatine-coated slides and mounted with Fluoromount-G with DAPI. Images were taken with a Zeiss LSM880 confocal microscope using the LSM image acquisition software Zen lite.

### Cell counting

The right brain hemisphere of each mouse was sliced in 50 μm coronal sections that were sequentially collected in series containing one every 8^th^ slice. Randomly chosen series of sections were used for immunohistochemistry against GFP and GFAP as described above. Due to the low numbers of M_4_^+^ astrocytes in the brain, we counted all visible cells positive for both GFP and GFAP within the indicated brain regions in every section of the series under a Zeiss microscope. DAPI images of each section were taken and the area of each region of interest was traced using the freehand drawing tool of ImageJ Fiji. The determined area was then multiplied by the thickness of the section to calculate the reference volume. The density of M_4_^+^ astrocytes (number of cells/mm^3^) was estimated by dividing the total number of positive cells by the reference volume of the corresponding region.

The quantification of M_4_^+^ astrocytes in the arbor vitae of the cerebellum was estimated by unbiased stereology using the physical dissector method adapted to confocal microscopy, as previously described (71). One in eight coronal sections of the cerebellum (501μm) were immunostained for GFP and GFAP as before. 9-12 confocal counting stacks, acquired with a LSM880 Zeiss microscope, were randomly selected from the whole arbor vitae of the cerebellum. The counting frame size was 1501×1150 μm with a thickness of 10.5 μm and a Z-interval of 1.5 μm. The mean cell density of M_4_^+^ astrocytes was estimated with the counts of the double-labelled cells and the size of the dissector.

### Saturation [^3^H]-NMS binding assay

Striata from homozygous M_4_ KO, R26-MetRS* (WT) or heterozygous Chrm4^CRE^::MetRS* were collected and homogenised in ice-cold HEPES buffer (10 mM HEPES, 1 mM EGTA, 1 mM dithiothreitol) containing 10% sucrose and protease inhibitors. The lysate was first centrifuged at 1,000 x g for 10 min at 4°C, the pellet resuspended, re-homogenised and further centrifuged at 20,000 x g for 1 h at 4°C. The resulting pellet containing striatal membranes, was diluted and protein concentration calculated by the Bradford method. 22.5 μg of the striatal membrane preparation were incubated in binding buffer (50 mM HEPEs, 110 mM NaCl, 5.4 mM KCl, 1.8 mM CaCl_2_, 1 mM MgSO_4_, 25 mM glucose, 58 mM sucrose, pH 7.4) with increasing concentrations (0.3-5 nM) of the muscarinic receptor antagonist [^3^H]-N-methyl scopolamine ([^3^H]-NMS, Cat. #NET636001MC, PerkinElmer) for 1 h at 37°C. Nonspecific binding was determined by the addition of atropine (1 μM) during the incubation with [^3^H]-NMS. Binding reactions were terminated by rapid filtration over UniFilter GF/C plates (PerkinElmer) pre-soaked in binding buffer, followed by three rapid washes with ice-cold 0.9% NaCl using a filter plate system (Brandel). The plates were then air-dried and 50 μL Microscint-20 liquid scintillation fluid (PerkinElmer) was added to each well. Membrane bound radioactivity was then counted in a Topcount liquid scintillation counter (PerkinElmer). The specific bound counts (dpm) were expressed as fmol/mg protein.

### Quantitative RT-PCR

Striata or cerebella from homozygous M_4_ KO and R26-MetRS* (WT) mice (10-12 weeks old) were collected and total RNA was isolated using a RNeasy Plus Mini Kit (Cat. #74134, Qiagen), following the manufacturer’s instructions. 1 µg total RNA was treated with DNase I (Cat. #79254, Qiagen) and used as template for cDNA synthesis with M-MLV reverse transcriptase (Cat. #28025013, Invitrogen), according to the manufacturer’s recommendations. Real-time PCR was performed with probes for Chrm4 (forward 5’-GGAGAAGAAGGCCAAGACTCTGG and reverse 5’-GGCAGTCACACATTCACTGCCTG) and GAPDH as control gene (forward 5’-CGGATTTGGCCGTATTGGG and reverse 5’-CTCGCTCCTGGAAGATGG), 2.4 μl of the cDNA sample as a template and Fast SYBR Green Master Mix (Cat. #4385612, Applied Biosystems) on a Quantstudio 5 Real Time PCR system (ThermoFisher Scientific). The cycling conditions were 50°C for 2 min, 95°C for 10 min, followed by 45 cycles of 95°C for 15 s and 60°C for 1 min. Melting curve analyses were performed for all real-time PCR runs to confirm amplification of only one product and to eliminate false positive results. Quantitative analysis was performed on Quantstudio software (ThermoFisher Scientific), which calculated the threshold cycle (Ct) values. Expression of Chrm4 was defined relative to GAPDH using the 2−ΔΔCt method.

### BONCAT purification for LC-MS/MS analysis

Cerebellar primary astrocytes from Chrm4^CRE^::MetRS* mice grown onto 10 cm dishes as before, were incubated with 4 mM ANL in combination with vehicle, oxotremorine (10 µM) or oxotremorine in combination with VU6028418 (5 µM) for 24 h at 37°C. Cells were lysed in PBS with 0.5% SDS, 0.2% Triton X-100 and proteinase inhibitors, briefly sonicated with 2 pulses of 15 s at an amplitude of 40% (∼50 W) and centrifuged at 10,000 x g for 10 min at 4°C. Supernatants were collected and protein concentration calculated with a BCA assay kit. 500 µg of protein were heated at 65°C for 5 min followed by incubation with 50 mM iodoacetamide for 2 h at room temperature in the dark. Bio-orthogonal non-canonical amino acid tagging (BONCAT) was performed using 150 µM Biotin-PEG4-alkyne (Cat. #764213, Sigma-Aldrich), 0.5 mM CuSO_4_, 1 mM THPTA and 5 mM sodium ascorbate overnight at 4°C. Buffer exchange was done with PD SpinTrap G-25 columns (Cat. #28-9180-04, Cytvia) and biotinylation of ANL-labelled proteins was confirmed by Western Blot. Ultralink avidin beads (Cat. #53146, Pierce) were used to purify clicked proteins. After overnight incubation at 4°C, beads were loaded into micro-spin columns (Cat. #89879, Pierce) and washed three times with PBS and three washes with ultrapure water. Proteins were eluted with 125 µl of 0.1% Trifluoroacetic acid (TFA) followed by 125 µl of 0.1% TFA with 50% acetonitrile (ACN). Eluted proteins were digested with 4 µg Trypsin/Lys-C (Cat. #V5073, Promega) in S-Trap midi spin columns (Profiti) according to the manufacturer’s instructions.

Dried peptides were resuspended in 5 % acetonitrile with 0.5 % formic acid for nanoflow liquid chromatography–electrospray ionisation-tandem mass spectrometry (nLC-ESI-MS/MS) analysis. The samples were loaded onto a C_18_ Acclaim PepMap trapping column (ThermoFisher Scientific) using an UltiMate3000 RSLCnano System (ThermoFisher Scientific). Peptide separation was performed on a Pepmap C_18_ reversed phase column (2 μm, 75 μm × 50 cm, pore size 100 Å, Cat. #164540, ThermoFisher Scientific) by a gradient formed by mobile phase A (0.1% formic acid) and B (80% ACN, 0.08% formic acid), with increasing concentrations of solvent B running from 4% for 10 min, 60% for 170 min to 99% for 15 min at a fixed solvent flow rate of 0.3 µl/min. Eluted peptides were electrosprayed into an Orbitrap Elite MS (ThermoFisher Scientific) using a NanoMate Triversa (Advion Biosciences) ion source at 1.7 kV. The mass spectrometer was operated using full-scan MS spectra (m/z 375–1,500) at a resolution of 120,000 RP, followed by top speed (3s) high energy collision dissociation (HCD) fragmentation detection of the top precursor ions. Ion fragmentation was performed by top speed (3s) high energy collision dissociation (HCD) and the top precursor ions were detected at a resolution of 60,000 in MS2 spectra. HCD energy and isolation width was adjusted depending on charge of the precursor ions (a width of 1.2 m/z with 40% HCD for 2^+^ ions, 0.7 m/z with 38% for 3^+^, and 0.5 m/z with 38% HCD for 4^+^-6^+^). The resulting MS/MS spectra were analysed for protein identification using Proteome Discoverer software 2.4 (ThermoFisher Scientific) and searched against the murine UniProtKB/Swiss-Prot database using Mascot software (Matrix Science) with peptide tolerance set to 10 ppm and fragment ion mass tolerance of 0.3 Da. The carbamidomethylation of cysteine was programmed as fixed modification, while the oxidation of methionine, the deamidation of asparagine and glutamine, the replacement of methionine for ANL-PEG-biotin and the iodination of tyrosine were set as variables. Proteins present in at least 2 independent experiments were considered for GO enrichment analysis to identify overrepresented biological processes using PANTHER (72) with a Fisher’s exact test and a cutoff of FDR ≤0.05.

### Statistical analysis

All data were expressed as mean ± SEM of at least 3 independent experiments. Statistical significance was determined by two-tailed Student’s *t* test, one-way ANOVA with Tukey’s post hoc correction or two-way ANOVA followed by a Tukey’s or Bonferroni’s post hoc test, as indicated. For ERK_1/2_ phosphorylation assays the concentration–response data were plotted on a log axis, fitted to a three-parameter sigmoidal curve and extra sum-of-squares F test applied to determine whether differences in pEC_50_ existed. The Kaplan-Meier plot survival analyses were compared using the Gehan-Breslow-Wilcoxon test. Significance was considered a *p* value smaller than 0.05 (\**p* < 0.05, \*\**p* < 0.01, \*\*\**p* < 0.001). Statistical analysis and graph plotting was performed using GraphPad Prism software.

## Supporting information

Supplementary uncropped immunoblots

## Abbreviations

CNS: central nervous system
ERK_1/2_: extracellular signal-regulated kinases ½
GFAP: glial fibrillary acidic protein
GFP: green fluorescent protein
eGFP: enhanced green fluorescent protein
GPCR: G protein-coupled receptor
mAChR: muscarinic acetylcholine receptor
NBH: normal brain homogenate
PAM: positive allosteric modulator
pEC: half maximal response
Prp^Sc^: scrapie prion protein
RML: rocky mountain laboratory
w.p.i: weeks post-inoculation.

## Declarations

### Ethics approval

All animal experiments were conducted in accordance with the UK Home Office Animal Procedures Act (1986) and all procedures were approved by the University of Glasgow Animal Welfare and Ethical Review Board.

### Consent for publication

Not applicable

### Availability of data and materials

Source data are provided with this paper.

## Competing Interests

The authors declare no competing interests.

## Funding

This work was supported by grants from the Wellcome Trust (Collaborative Award 201529/Z/16/Z and Discovery Award 307319/Z/23/Z).

## Author’s Contributions

GST conducted the experiments, developed the study design and wrote the paper, ABT lead the study design and wrote the paper, GM contributed to the study design and writing of the paper, CM managed the mouse colonies and breeding and helped to conduct the animal experiments, CWL contributed with the muscarinic antagonist VU6028418 and the M_4_ mAChR positive allosteric modulator VU0467154 and CKJ contributed with pharmacokinetic, pharmacology and metabolism data of these ligands.

## Acknowledgments

The authors would like to thank the staff of the University of Glasgow Biomedical Services, genOway for generation of the muscarinic receptor genetically modified mice. The authors would also like to acknowledge the assistance of Dr. Karen Fairlie-Clarke to this work.

## Notes

### Competing Interest Statement

The authors have declared no competing interest.

### Summary of Updates

This version of the manuscript has been revised to incorporate several updates aimed at improving clarity, organisation, and data transparency. The title and abstract have been shortened to provide a more concise summary of the study. The introduction has been restructured to enhance readability and strengthen the logical flow of the narrative.Textual refinements have been applied throughout the manuscript to improve precision and consistency. A comprehensive Supplementary Data file has been added, including uncropped western blot images and an Excel file containing all numerical data used in figure quantifications. New experimental data have been included in Figure 5e and Supplementary Figure 5f to expand the scope of the results. In addition, certain figures have been reorganized within the manuscript to improve alignment with the main text and overall presentation. These updates collectively aim to ensure clarity, transparency, and completeness of the scientific record.

## References

1. Kaul I, Sawchak S, Correll CU, Kakar R, Breier A, Zhu H, et al. Efficacy and safety of the muscarinic receptor agonist KarXT (xanomeline-trospium) in schizophrenia (EMERGENT-2) in the USA: results from a randomised, double-blind, placebo-controlled, flexible-dose phase 3 trial. Lancet. 2024;403(10422):160–70.

2. Kaul I, Sawchak S, Walling DP, Tamminga CA, Breier A, Zhu H, et al. Efficacy and Safety of Xanomeline-Trospium Chloride in Schizophrenia: A Randomized Clinical Trial. JAMA Psychiatry. 2024;81(8):749–56.

3. Mei F, Lehmann-Horn K, Shen YA, Rankin KA, Stebbins KJ, Lorrain DS, et al. Accelerated remyelination during inflammatory demyelination prevents axonal loss and improves functional recovery. Elife. 2016;5.

4. Wang F, Yang YJ, Yang N, Chen XJ, Huang NX, Zhang J, et al. Enhancing Oligodendrocyte Myelination Rescues Synaptic Loss and Improves Functional Recovery after Chronic Hypoxia. Neuron. 2018;99(4):689–701 e5.

5. Wang F, Ren SY, Chen JF, Liu K, Li RX, Li ZF, et al. Myelin degeneration and diminished myelin renewal contribute to age-related deficits in memory. Nat Neurosci. 2020;23(4):481–6.

6. Chen JF, Liu K, Hu B, Li RR, Xin W, Chen H, et al. Enhancing myelin renewal reverses cognitive dysfunction in a murine model of Alzheimer’s disease. Neuron. 2021;109(14):2292–307 e5.

7. Dwomoh L, Rossi M, Scarpa M, Khajehali E, Molloy C, Herzyk P, et al. M(1) muscarinic receptor activation reduces the molecular pathology and slows the progression of prion-mediated neurodegenerative disease. Sci Signal. 2022;15(760):eabm3720.

8. Abd-Elrahman KS, Colson TL, Sarasija S, Ferguson SSG. A M1 muscarinic acetylcholine receptor-specific positive allosteric modulator VU0486846 reduces neurogliosis in female Alzheimer’s mice. Biomed Pharmacother. 2024;173:116388.

9. Pannell M, Meier MA, Szulzewsky F, Matyash V, Endres M, Kronenberg G, et al. The subpopulation of microglia expressing functional muscarinic acetylcholine receptors expands in stroke and Alzheimer’s disease. Brain Struct Funct. 2016;221(2):1157–72.

10. Andre C, Dos Santos G, Koulakoff A. Muscarinic receptor profiles of mouse brain astrocytes in culture vary with their tissue of origin but differ from those of neurons. Eur J Neurosci. 1994;6(11):1702–9.

11. Endo F, Kasai A, Soto JS, Yu X, Qu Z, Hashimoto H, et al. Molecular basis of astrocyte diversity and morphology across the CNS in health and disease. Science. 2022;378(6619):eadc9020.

12. Zhang Y, Chen K, Sloan SA, Bennett ML, Scholze AR, O’Keeffe S, et al. An RNA-sequencing transcriptome and splicing database of glia, neurons, and vascular cells of the cerebral cortex. J Neurosci. 2014;34(36):11929–47.

13. Yao Z, van Velthoven CTJ, Kunst M, Zhang M, McMillen D, Lee C, et al. A high-resolution transcriptomic and spatial atlas of cell types in the whole mouse brain. Nature. 2023;624(7991):317–32.

14. Alvarez-Castelao B, Schanzenbacher CT, Hanus C, Glock C, Tom Dieck S, Dorrbaum AR, et al. Cell-type-specific metabolic labeling of nascent proteomes in vivo. Nat Biotechnol. 2017;35(12):1196–201.

15. Sjostedt E, Zhong W, Fagerberg L, Karlsson M, Mitsios N, Adori C, et al. An atlas of the protein-coding genes in the human, pig, and mouse brain. Science. 2020;367(6482).

16. Buckley NJ, Bonner TI, Brann MR. Localization of a family of muscarinic receptor mRNAs in rat brain. J Neurosci. 1988;8(12):4646–52.

17. Heintz N. Gene expression nervous system atlas (GENSAT). Nat Neurosci. 2004;7(5):483.

18. Vilaro MT, Wiederhold KH, Palacios JM, Mengod G. Muscarinic cholinergic receptors in the rat caudate-putamen and olfactory tubercle belong predominantly to the m4 class: in situ hybridization and receptor autoradiography evidence. Neuroscience. 1991;40(1):159–67.

19. Arenander AT, de Vellis J, Herschman HR. Induction of c-fos and TIS genes in cultured rat astrocytes by neurotransmitters. J Neurosci Res. 1989;24(1):107–14.

20. Bubser M, Bridges TM, Dencker D, Gould RW, Grannan M, Noetzel MJ, et al. Selective activation of M4 muscarinic acetylcholine receptors reverses MK-801-induced behavioral impairments and enhances associative learning in rodents. ACS Chem Neurosci. 2014;5(10):920–42.

21. Spock M, Carter TR, Bollinger KA, Han C, Baker LA, Rodriguez AL, et al. Discovery of VU6028418: A Highly Selective and Orally Bioavailable M(4) Muscarinic Acetylcholine Receptor Antagonist. ACS Med Chem Lett. 2021;12(8):1342–9.

22. Tsai HH, Li H, Fuentealba LC, Molofsky AV, Taveira-Marques R, Zhuang H, et al. Regional astrocyte allocation regulates CNS synaptogenesis and repair. Science. 2012;337(6092):358–62.

23. Cerrato V. Cerebellar Astrocytes: Much More Than Passive Bystanders In Ataxia Pathophysiology. J Clin Med. 2020;9(3).

24. Gould RW, Grannan MD, Gunter BW, Ball J, Bubser M, Bridges TM, et al. Cognitive enhancement and antipsychotic-like activity following repeated dosing with the selective M(4) PAM VU0467154. Neuropharmacology. 2018;128:492–502.

25. Foster DJ, Wilson JM, Remke DH, Mahmood MS, Uddin MJ, Wess J, et al. Antipsychotic-like Effects of M4 Positive Allosteric Modulators Are Mediated by CB2 Receptor-Dependent Inhibition of Dopamine Release. Neuron. 2016;91(6):1244–52.

26. Pancani T, Bolarinwa C, Smith Y, Lindsley CW, Conn PJ, Xiang Z. M4 mAChR-mediated modulation of glutamatergic transmission at corticostriatal synapses. ACS Chem Neurosci. 2014;5(4):318–24.

27. Moehle MS, Pancani T, Byun N, Yohn SE, Wilson GH, 3rd, Dickerson JW, et al. Cholinergic Projections to the Substantia Nigra Pars Reticulata Inhibit Dopamine Modulation of Basal Ganglia through the M(4) Muscarinic Receptor. Neuron. 2017;96(6):1358–72 e4.

28. Jeon J, Dencker D, Wortwein G, Woldbye DP, Cui Y, Davis AA, et al. A subpopulation of neuronal M4 muscarinic acetylcholine receptors plays a critical role in modulating dopamine-dependent behaviors. J Neurosci. 2010;30(6):2396–405.

29. Zimmer TS, Orr AL, Orr AG. Astrocytes in selective vulnerability to neurodegenerative disease. Trends Neurosci. 2024;47(4):289–302.

30. Endo F. Deciphering the spectrum of astrocyte diversity: Insights into molecular, morphological, and functional dimensions in health and neurodegenerative diseases. Neurosci Res. 2024.

31. Moulson AJ, Squair JW, Franklin RJM, Tetzlaff W, Assinck P. Diversity of Reactive Astrogliosis in CNS Pathology: Heterogeneity or Plasticity? Front Cell Neurosci. 2021;15:703810.

32. Tahir W, Thapa S, Schatzl HM. Astrocyte in prion disease: a double-edged sword. Neural Regen Res. 2022;17(8):1659–65.

33. Hartmann K, Sepulveda-Falla D, Rose IVL, Madore C, Muth C, Matschke J, et al. Complement 3(+)-astrocytes are highly abundant in prion diseases, but their abolishment led to an accelerated disease course and early dysregulation of microglia. Acta Neuropathol Commun. 2019;7(1):83.

34. Akhtar S, Wenborn A, Brandner S, Collinge J, Lloyd SE. Sex effects in mouse prion disease incubation time. PLoS One. 2011;6(12):e28741.

35. Liu Y, Guo J, Matoga M, Korotkova M, Jakobsson PJ, Aguzzi A. NG2 glia protect against prion neurotoxicity by inhibiting microglia-to-neuron prostaglandin E2 signaling. Nat Neurosci. 2024;27(8):1534–44.

36. Denouel A, Brandel JP, Seilhean D, Laplanche JL, Elbaz A, Haik S. The role of environmental factors on sporadic Creutzfeldt-Jakob disease mortality: evidence from an age-period-cohort analysis. Eur J Epidemiol. 2023;38(7):757–64.

37. Myslivecek J, Farar V, Valuskova P. M(4) muscarinic receptors and locomotor activity regulation. Physiol Res. 2017;66(Suppl 4):S443–S55.

38. Chambers NE, Millett M, Moehle MS. The muscarinic M4 acetylcholine receptor exacerbates symptoms of movement disorders. Biochem Soc Trans. 2023;51(2):691–702.

39. Nunes EJ, Addy NA, Conn PJ, Foster DJ. Targeting the Actions of Muscarinic Receptors on Dopamine Systems: New Strategies for Treating Neuropsychiatric Disorders. Annu Rev Pharmacol Toxicol. 2024;64:277–89.

40. Camandola S. Astrocytes, emerging stars of energy homeostasis. Cell Stress. 2018;2(10):246–52.

41. Gonzalez-Garcia I, Garcia-Caceres C. Hypothalamic Astrocytes as a Specialized and Responsive Cell Population in Obesity. Int J Mol Sci. 2021;22(12).

42. Yang L, Qi Y, Yang Y. Astrocytes control food intake by inhibiting AGRP neuron activity via adenosine A1 receptors. Cell Rep. 2015;11(5):798–807.

43. Chen N, Sugihara H, Kim J, Fu Z, Barak B, Sur M, et al. Direct modulation of GFAP-expressing glia in the arcuate nucleus bi-directionally regulates feeding. Elife. 2016;5.

44. Herrera Moro Chao D, Kirchner MK, Pham C, Foppen E, Denis RGP, Castel J, et al. Hypothalamic astrocytes control systemic glucose metabolism and energy balance. Cell Metab. 2022;34(10):1532–47 e6.

45. MacDonald AJ, Holmes FE, Beall C, Pickering AE, Ellacott KLJ. Regulation of food intake by astrocytes in the brainstem dorsal vagal complex. Glia. 2020;68(6):1241–54.

46. Clyburn C, Carson KE, Smith CR, Travagli RA, Browning KN. Brainstem astrocytes control homeostatic regulation of caloric intake. J Physiol. 2023;601(4):801–29.

47. Cai P, Huang SN, Lin ZH, Wang Z, Liu RF, Xiao WH, et al. Regulation of wakefulness by astrocytes in the lateral hypothalamus. Neuropharmacology. 2022;221:109275.

48. Brancaccio M, Edwards MD, Patton AP, Smyllie NJ, Chesham JE, Maywood ES, et al. Cell-autonomous clock of astrocytes drives circadian behavior in mammals. Science. 2019;363(6423):187–92.

49. Nimmerjahn A, Mukamel EA, Schnitzer MJ. Motor behavior activates Bergmann glial networks. Neuron. 2009;62(3):400–12.

50. Paukert M, Agarwal A, Cha J, Doze VA, Kang JU, Bergles DE. Norepinephrine controls astroglial responsiveness to local circuit activity. Neuron. 2014;82(6):1263–70.

51. Corkrum M, Covelo A, Lines J, Bellocchio L, Pisansky M, Loke K, et al. Dopamine-Evoked Synaptic Regulation in the Nucleus Accumbens Requires Astrocyte Activity. Neuron. 2020;105(6):1036–47 e5.

52. Schweinhuber SK, Messerschmidt T, Hansch R, Korte M, Rothkegel M. Profilin isoforms modulate astrocytic morphology and the motility of astrocytic processes. PLoS One. 2015;10(1):e0117244.

53. Lee SJ, Seo BR, Koh JY. Metallothionein-3 modulates the amyloid beta endocytosis of astrocytes through its effects on actin polymerization. Mol Brain. 2015;8(1):84.

54. Zhou J, Zhang L, Peng J, Zhang X, Zhang F, Wu Y, et al. Astrocytic LRP1 enables mitochondria transfer to neurons and mitigates brain ischemic stroke by suppressing ARF1 lactylation. Cell Metab. 2024;36(9):2054–68 e14.

55. Romeo R, Glotzbach K, Scheller A, Faissner A. Deletion of LRP1 From Astrocytes Modifies Neuronal Network Activity in an in vitro Model of the Tripartite Synapse. Front Cell Neurosci. 2020;14:567253.

56. Gogliotti RG, Fisher NM, Stansley BJ, Jones CK, Lindsley CW, Conn PJ, et al. Total RNA Sequencing of Rett Syndrome Autopsy Samples Identifies the M(4) Muscarinic Receptor as a Novel Therapeutic Target. J Pharmacol Exp Ther. 2018;365(2):291–300.

57. Achilly NP, He LJ, Kim OA, Ohmae S, Wojaczynski GJ, Lin T, et al. Deleting Mecp2 from the cerebellum rather than its neuronal subtypes causes a delay in motor learning in mice. Elife. 2021;10.

58. Patani R, Hardingham GE, Liddelow SA. Functional roles of reactive astrocytes in neuroinflammation and neurodegeneration. Nat Rev Neurol. 2023;19(7):395–409.

59. Makarava N, Chang JC, Kushwaha R, Baskakov IV. Region-Specific Response of Astrocytes to Prion Infection. Front Neurosci. 2019;13:1048.

60. Makarava N, Chang JC, Molesworth K, Baskakov IV. Region-specific glial homeostatic signature in prion diseases is replaced by a uniform neuroinflammation signature, common for brain regions and prion strains with different cell tropism. Neurobiol Dis. 2020;137:104783.

61. Carroll JA, Striebel JF, Rangel A, Woods T, Phillips K, Peterson KE, et al. Prion Strain Differences in Accumulation of PrPSc on Neurons and Glia Are Associated with Similar Expression Profiles of Neuroinflammatory Genes: Comparison of Three Prion Strains. PLoS Pathog. 2016;12(4):e1005551.

62. Munoz-Castro C, Serrano-Pozo A. Astrocyte-Neuron Interactions in Alzheimer’s Disease. Adv Neurobiol. 2024;39:345–82.

63. Carter SF, Herholz K, Rosa-Neto P, Pellerin L, Nordberg A, Zimmer ER. Astrocyte Biomarkers in Alzheimer’s Disease. Trends Mol Med. 2019;25(2):77–95.

64. Makarava N, Mychko O, Chang JC, Molesworth K, Baskakov IV. The degree of astrocyte activation is predictive of the incubation time to prion disease. Acta Neuropathol Commun. 2021;9(1):87.

65. Gomeza J, Zhang L, Kostenis E, Felder CC, Bymaster FP, Brodkin J, et al. Generation and pharmacological analysis of M2 and M4 muscarinic receptor knockout mice. Life Sci. 2001;68(22-23):2457–66.

66. Guyenet SJ, Furrer SA, Damian VM, Baughan TD, La Spada AR, Garden GA. A simple composite phenotype scoring system for evaluating mouse models of cerebellar ataxia. J Vis Exp. 2010(39).

67. Mallucci G, Dickinson A, Linehan J, Klohn PC, Brandner S, Collinge J. Depleting neuronal PrP in prion infection prevents disease and reverses spongiosis. Science. 2003;302(5646):871–4.

68. Scarpa M, Molloy C, Jenkins L, Strellis B, Budgett RF, Hesse S, et al. Biased M1 muscarinic receptor mutant mice show accelerated progression of prion neurodegenerative disease. Proc Natl Acad Sci U S A. 2021;118(50).

69. Mirabile I, Jat PS, Brandner S, Collinge J. Identification of clinical target areas in the brainstem of prion-infected mice. Neuropathol Appl Neurobiol. 2015;41(5):613–30.

70. Wolfes AC, Dean C. Culturing In Vivo-like Murine Astrocytes Using the Fast, Simple, and Inexpensive AWESAM Protocol. J Vis Exp. 2018(131).

71. Llorens-Martin M, Torres-Aleman I, Trejo JL. Pronounced individual variation in the response to the stimulatory action of exercise on immature hippocampal neurons. Hippocampus. 2006;16(5):480–90.

72. Thomas PD, Ebert D, Muruganujan A, Mushayahama T, Albou LP, Mi H. PANTHER: Making genome-scale phylogenetics accessible to all. Protein Sci. 2022;31(1):8–22.

