## Supplementary Figures for "Astrocytes expressing M_4_ muscarinic acetylcholine receptor regulate locomotion and survival in murine prion disease"

**This PDF file includes:**

Supplementary Fig. 1-6

### Supplementary Fig. 1

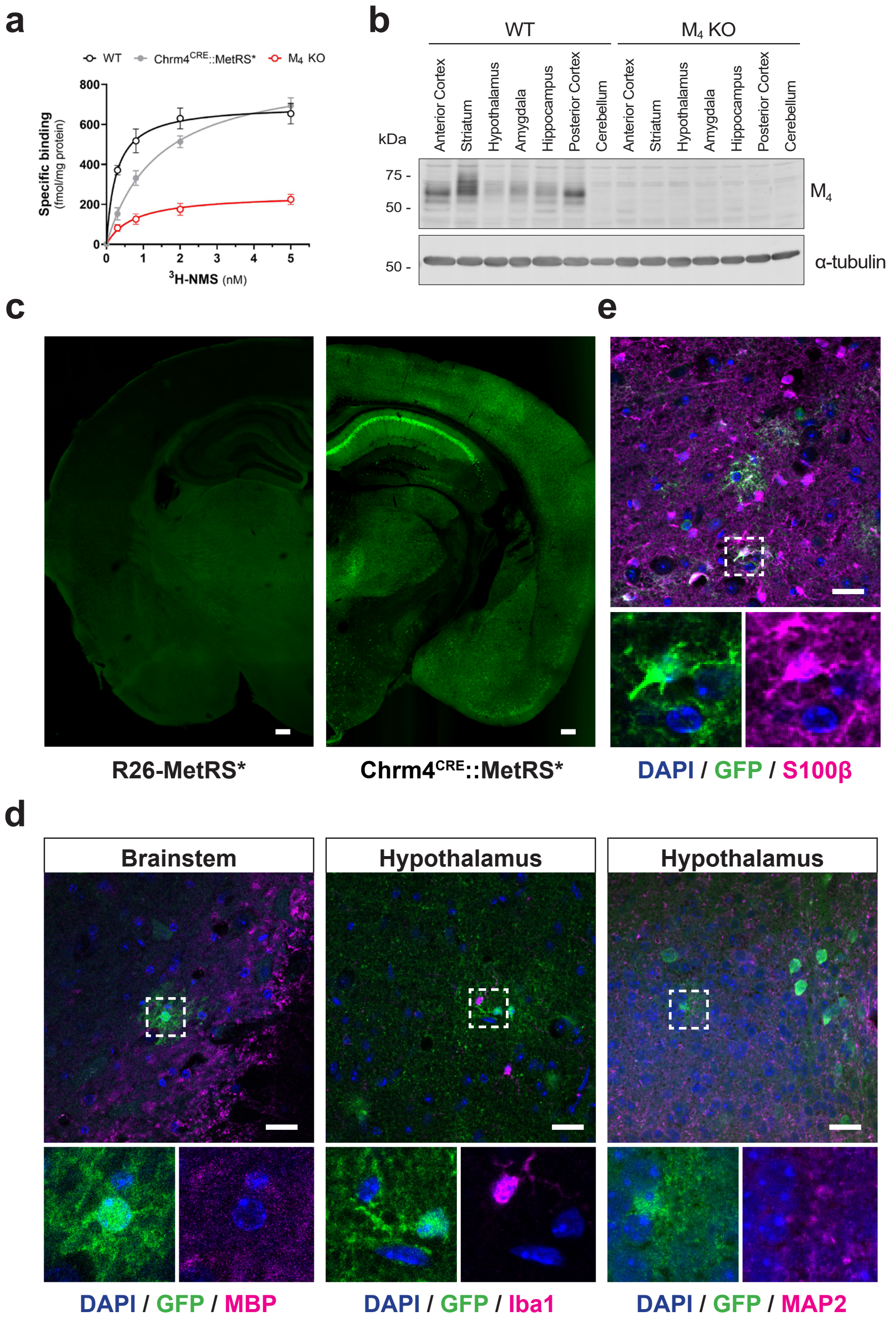


**Supplementary Fig. 1: Characterisation of M$_4$ mAChR expression and M$_4^+$ stellate cells in Chrm4$^{CRE}$::MetRS* mice. a**, [$^3$H]-N-methyl scopolamine (NMS) binding to striatal membranes prepared from wild type (WT), M$_4$ KO or Chrm4$^{CRE}$::MetRS* (heterozygous for both alleles) mice. Data are presented as mean ± SEM (n = 3). **b**, Immunoblot showing M$_4$ mAChR protein levels in different regions of the brain in WT and M$_4$ KO mice. α-tubulin was used as a loading control. **c**, Comparison of GFP staining (green) in brain coronal sections from the inducible R26-MetRS* (left) and Chrm4$^{CRE}$::MetRS*. **d**, Immunostaining analysis of GFP-labelled M$_4^+$ stellate cells (green) with markers for oligodendrocytes (MBP), microglia (Iba1) and neurons (MAP2) in the brainstem and hypothalamus of Chrm4$^{CRE}$::MetRS* mice. **e**, Co-localisation of GFP staining and the astrocyte marker S100β (magenta) in the hypothalamus and brainstem of Chrm4$^{CRE}$::MetRS* mice. Scale bar for panel **c** is 250 μm, and for **d** and **e**, 25 μm.

### Supplementary Fig. 2

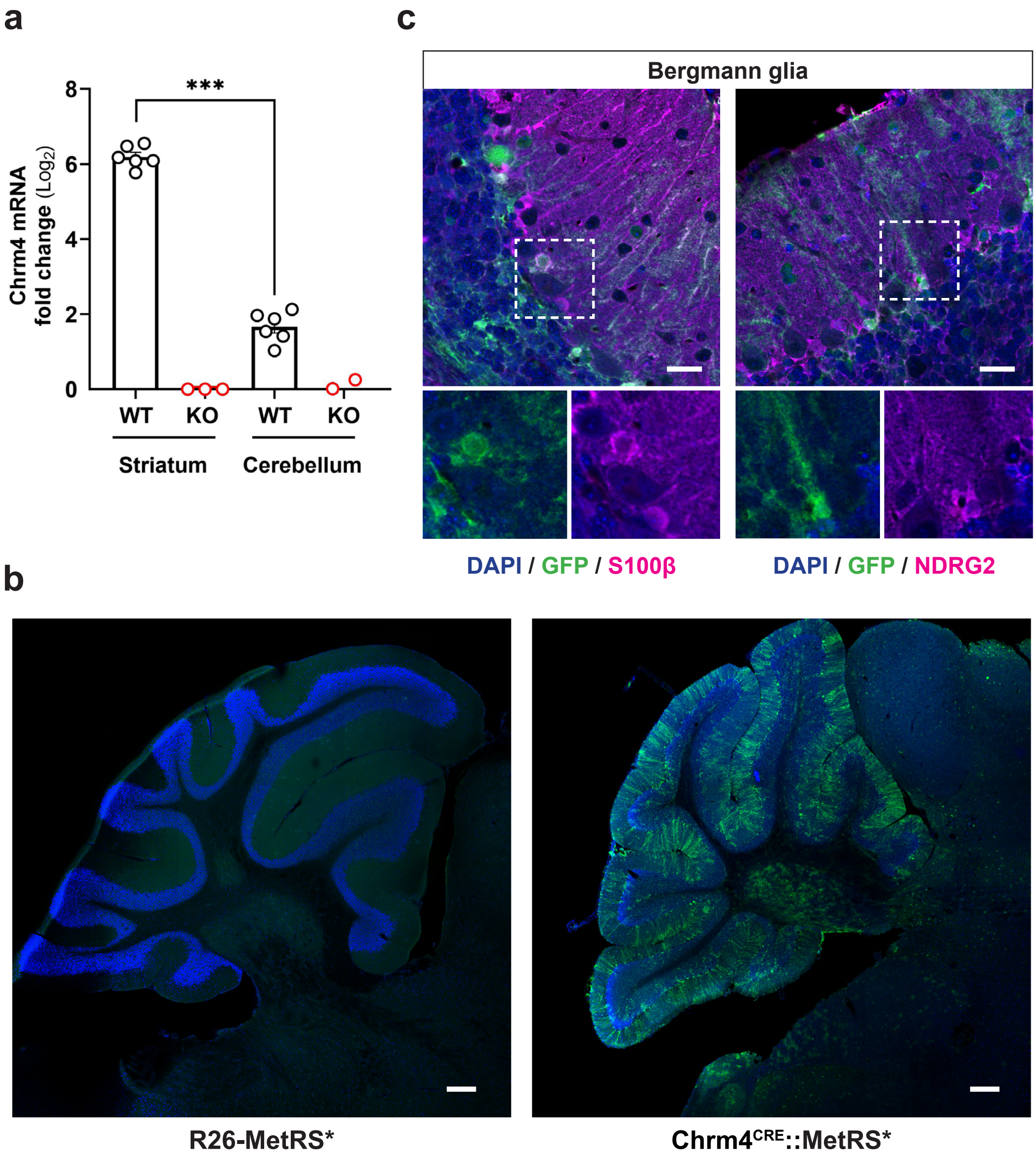


**Supplementary Fig. 2: Expression of $M_4$ mAChR in the cerebellum. a**, $M_4$ mAChR mRNA expression levels in striata and cerebella of WT mouse relative to values of $M_4$ KO in animals each tissue. Results are represented as $log_2$-fold change mean values ± SEM and statistical analysis was performed by one-way ANOVA followed by Tukey post hoc test (****p* < 0.001; n = 3 - 6). **b**, Expression pattern of GFP-immunostained cells in the cerebellum of the inducible R26-MetRS* (left) and $Chrm4^{CRE}$::MetRS* (right) mice. **c**, Co-localisation analysis between $M_4^+$ cells (green) with the astrocytic markers S100β (left) or NDRG2 (right, magenta) in Bergmann glia located in the cerebellar cortex of $Chrm4^{CRE}$::MetRS* mice. Scale bar for panel **b** is 250 µm, and for **c**, 25 µm.

### Supplementary Fig. 3

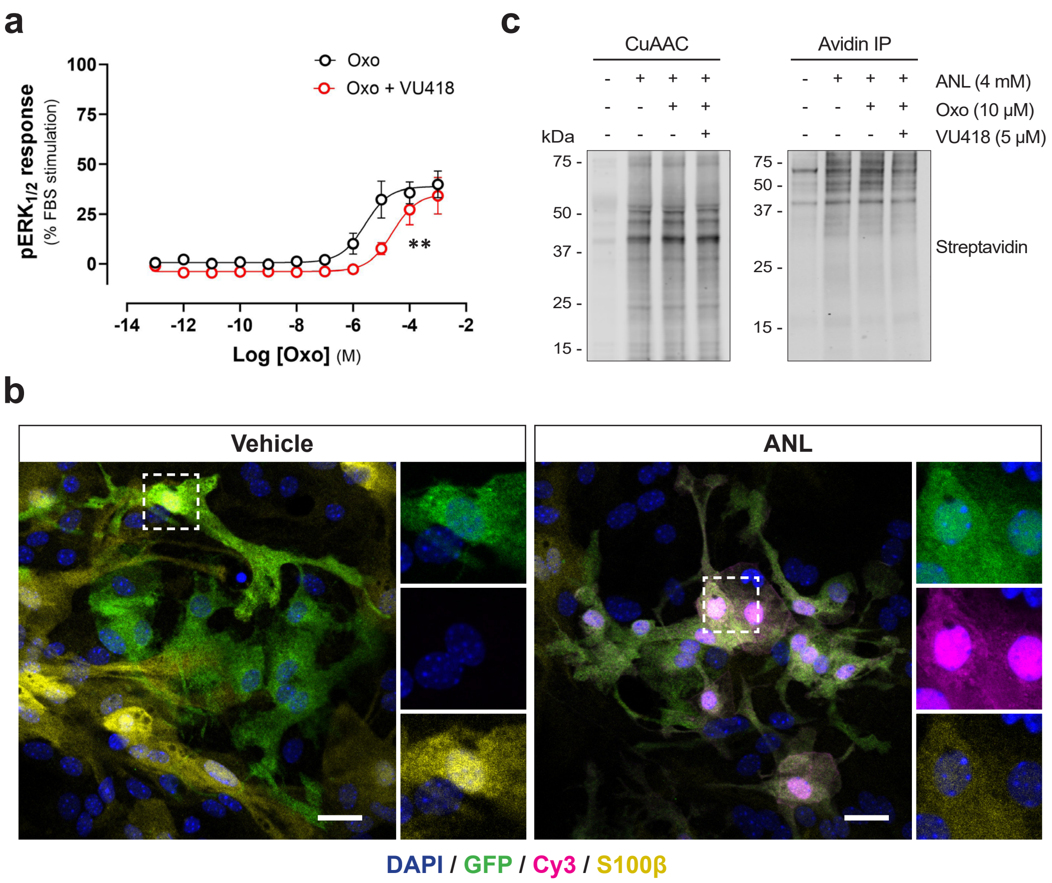


**Supplementary Fig. 3: $M_4$ mAChR-mediated activation in mouse cerebellar astrocytes.** **a**, $ERK_{1/2}$ phosphorylation assay in astrocyte cerebellar cultures from wild-type mice after stimulation with increasing concentrations of Oxo or Oxo in combination with the $M_4$ mAChR antagonist VU6028418 (VU418, 1 μM) (n = 3). Extra sum-of-squares F test was performed to evaluate differences in $pEC_{50}$ (**$p < 0.01$). **b**, Representative confocal images showing the selective labelling of Cy3-tagged NSPs (magenta) in $M_4^+$ astrocytes (green) after treatment with ANL (4 mM) in Chrm4$^{CRE}$::MetRS* cerebellar cultures which show staining for the astrocytic marker S100β (yellow). Scale bar is 25 μm. **c**, Immunoblot detection of biotin-labelled newly synthesized proteins (NSPs) from Chrm4$^{CRE}$::MetRS* cerebellar astrocyte cultures after treatment with ANL, in combination with Oxo or VU418. NSPs that incorporated ANL were biotinylated by copper-catalysed azide-alkyne cycloaddition (CuAAC) and later isolated by immunoprecipitation with avidin beads.

### Supplementary Fig. 4

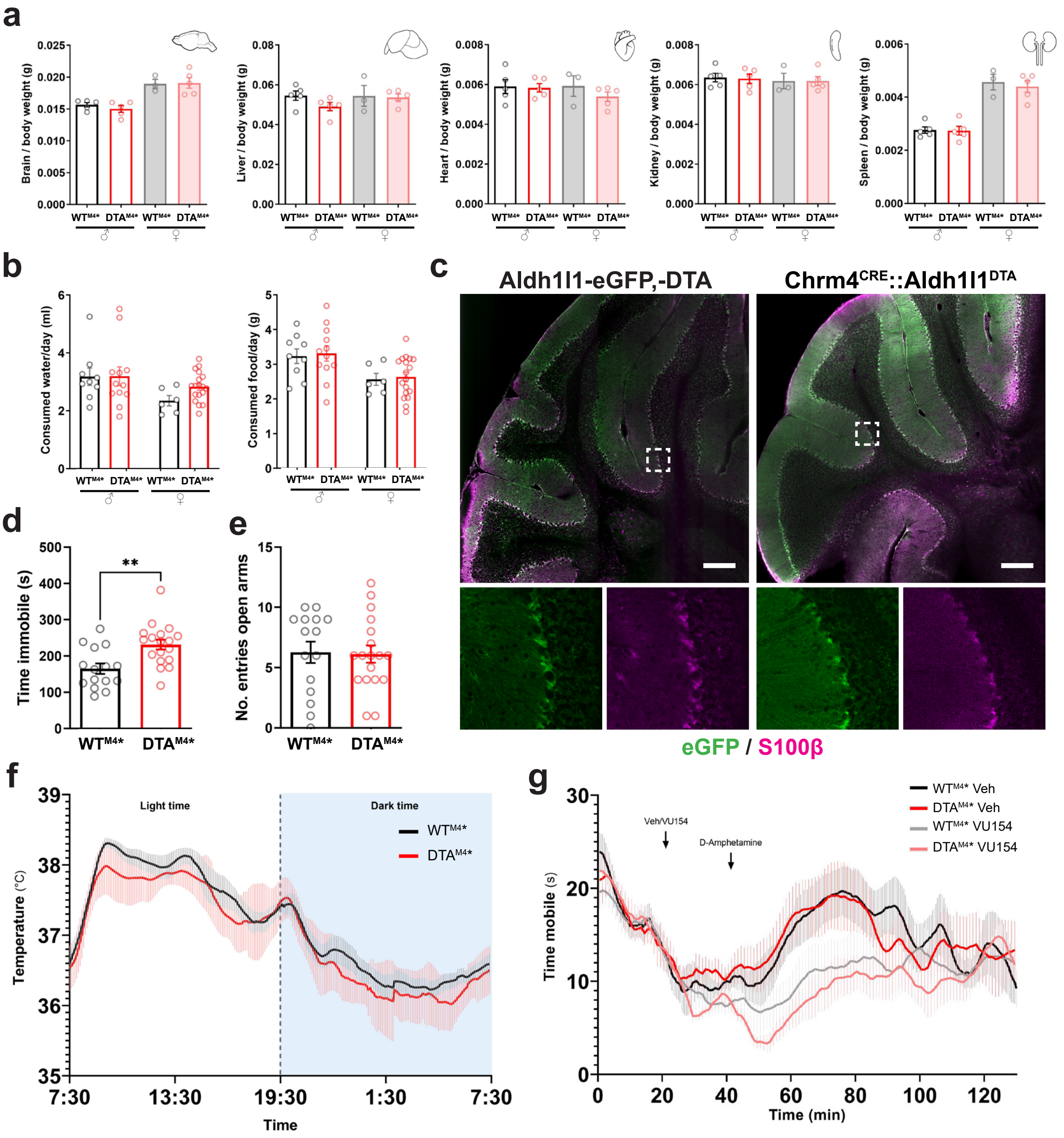


**Supplementary Fig. 4: Characterisation of targeted ablation of $M_4^+$ astrocytes in mice.** **a**, Average weight of brain, liver, heart, spleen and kidneys from DTA$^{M4*}$ and WT$^{M4*}$ male and female animals adjusted for body weight (n = 3 - 5). **b**, Comparison of the average daily food and water consumption per animal between DTA$^{M4*}$ and WT$^{M4*}$ mice (n = 6 - 18). **c**, Immunostaining analysis of the Bergmann glia distribution in DTA$^{M4*}$ mice (Chrm4$^{CRE}$::Aldh1l1$^{DTA}$, right) and the parental line (Aldh1l1-eGFP,-DTA, left). All detected S100β-labelled astrocytes (magenta) were co-stained with eGFP (green), expressed under the *Aldh1l1* promoter. Scale bar is 250 µm. **d**, Behavioural analysis of the immobility time during open field test of DTA$^{M4*}$ and WT$^{M4*}$ mice (n = 15 - 18). **e**, Measurement of the number of entries in the open arms of an elevated plus maze of DTA$^{M4*}$ and WT$^{M4*}$ mice (n = 15 - 18). **f**, Average daily abdominal temperature of DTA$^{M4*}$ and WT$^{M4*}$ mice measured by telemetry every five min over a period of 3 days (n = 8). **g**, Measurement of the mobility time after amphetamine-induced hyperlocomotion in DTA$^{M4*}$ and WT$^{M4*}$ mice treated with vehicle (Veh) or VU0467154 (VU154, 10 mg/kg). Data represents means ± SEM. Statistical significance in **d** was calculated by two-tailed unpaired *t* test (**$p$ < 0.01).

### Supplementary Fig. 5

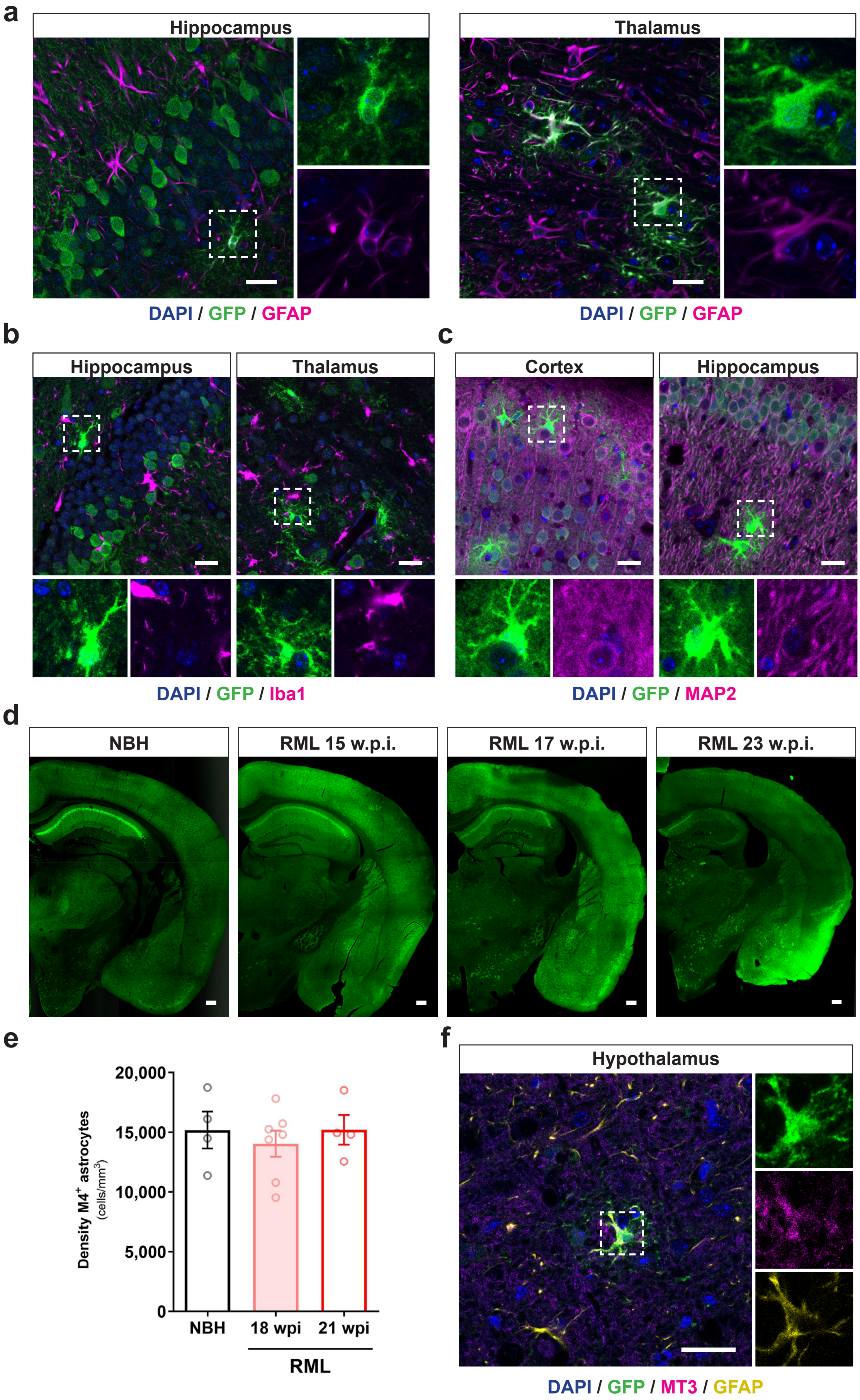

**Supplementary Fig. 5: Spreading of reactive $M_4^+$ astrocytes across diverse brain regions during prion disease. a**, Representative images showing co-localisation of GFP-labelled $M_4^+$ astrocytes with GFAP in the hippocampus and thalamus of 20 w.p.i. prion diseased Chrm4$^{CRE}$::MetRS* animals. **b**, Confocal images showing immunostaining of Iba1-labelled microglia (magenta) and $M_4^+$ cells (green) in the hippocampus and the thalamus of 20 w.p.i. prion diseased Chrm4$^{CRE}$::MetRS* mice. **c**, Immunodetection of $M_4^+$ astrocytes, not co-stained with the neuronal marker MAP2, in the cortex and hippocampus of RML-inoculated Chrm4$^{CRE}$::MetRS* mice (20 w.p.i). **d**, Representative brain coronal sections of Chrm4$^{CRE}$::MetRS* mice showing the increase in the number of $M_4^+$ astrocytes at different stages after inoculation with RML prion (15, 17 and 23 w.p.i.) compared to NBH control animals. **e**, Stereological counting of $M4^+$ astrocytes density in the cerebellar nuclei of mice inoculated with RML prion (18 and 21 w.p.i.) or NBH (n = 5-7). **f**, Representative image showing co-staining of $M_4^+$ cells (green) with metallothionein 3 (MT3, magenta) and GFAP (yellow) in the thalamus of 20 w.p.i. prion diseased Chrm4$^{CRE}$::MetRS* mice. Scale bars are 25 µm in **a**, **b**, **c** and **f** and 250 µm in **d**.

### Supplementary Fig. 6

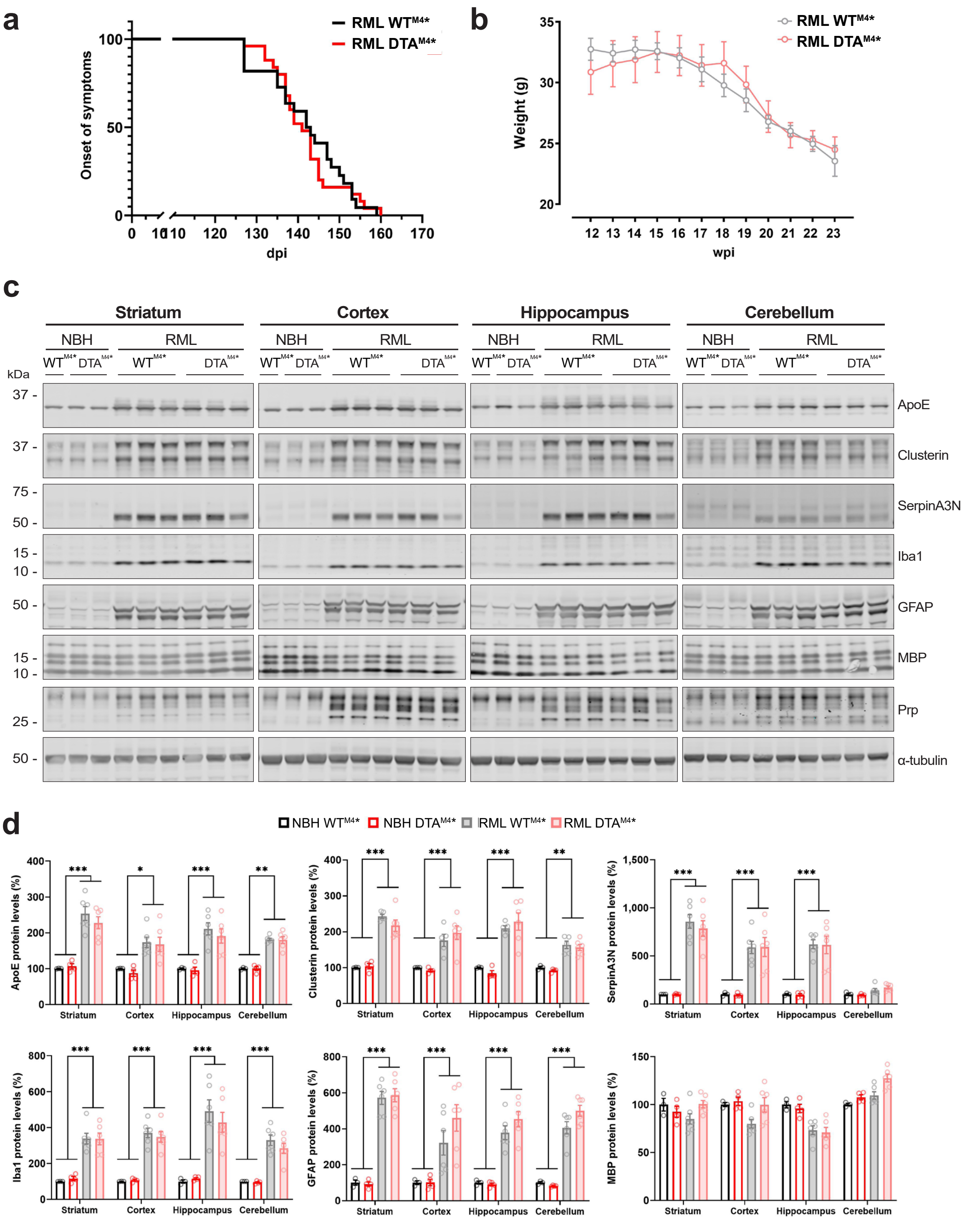


**Supplementary Fig. 6: Impact of $M_4^+$ astrocytes ablation in mice during prion disease.** **a**, Onset of at least two early indicator signs of prion disease (subdued behaviour, intermittent generalized tremor, erect penis, rigid tail, unsustained hunched posture, mild loss of coordination) in RML-inoculated mice with no $M_4^+$ astrocytes ($DTA^{M4*}$) or wild-type littermates ($WT^{M4*}$) (n = 22 - 25). **b**, Weight loss of prion-diseased $DTA^{M4*}$ and $WT^{M4*}$ male mice over time (n = 8 - 12). **c**, Western blot analysis of indicators of neurodegeneration (ApoE, cluterin, serpinA3N), markers of neuroinflammation (Iba1, GFAP, MBP) and Prp in the striatum, cortex, hippocampus and cerebellum from NBH and RML-inoculated $DTA^{M4*}$ and $WT^{M4*}$ mice. Samples were collected at terminal stage from animals with equivalent survival times. **d**, Densitometric analysis of the previously indicated proteins shown as means ± SEM of a ratio of α-tubulin expression relative to the expression obtained in the NBH $WT^{M4*}$animals (n = 4 - 6). Data were analysed using a two-way ANOVA with a Tukey post-hoc test (*$p < 0.05$, **$p < 0.01$, ***$p < 0.001$).
