## Supplementary uncropped immunoblots for "Astrocytes expressing M_4_ muscarinic acetylcholine receptor regulate locomotion and survival in murine prion disease"

### Source data

**Uncropped immunoblots related to Suppl. Fig. 6f and 6g**

* Precision Plus Protein Standards - BioRad 1610373

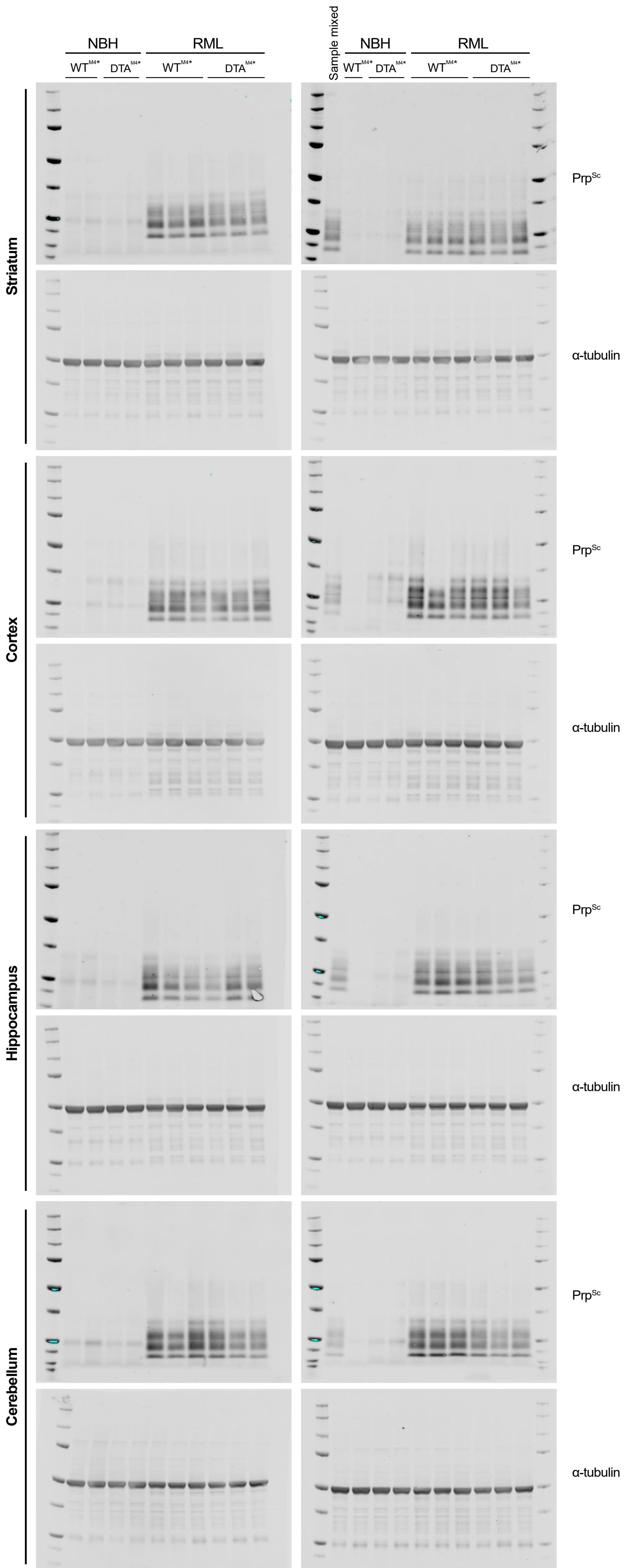

### Uncropped immunoblots related to Suppl. Fig. 1b

* Precision Plus Protein Standards -
BioRad 1610373

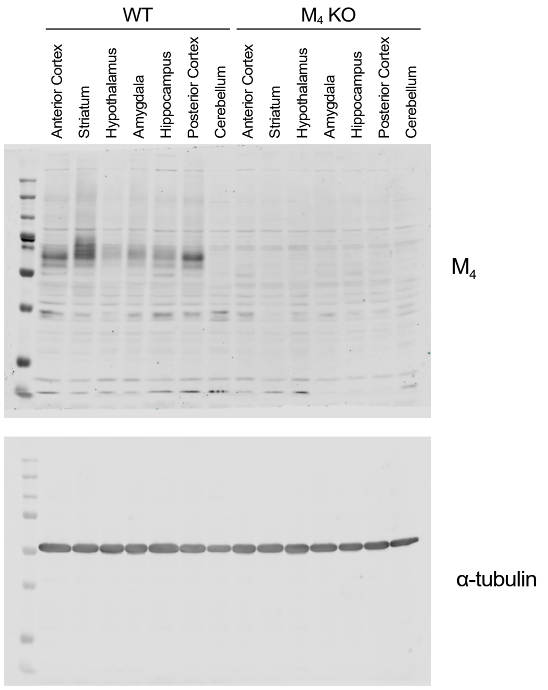


### Uncropped immunoblots related to Suppl. Fig. 3c

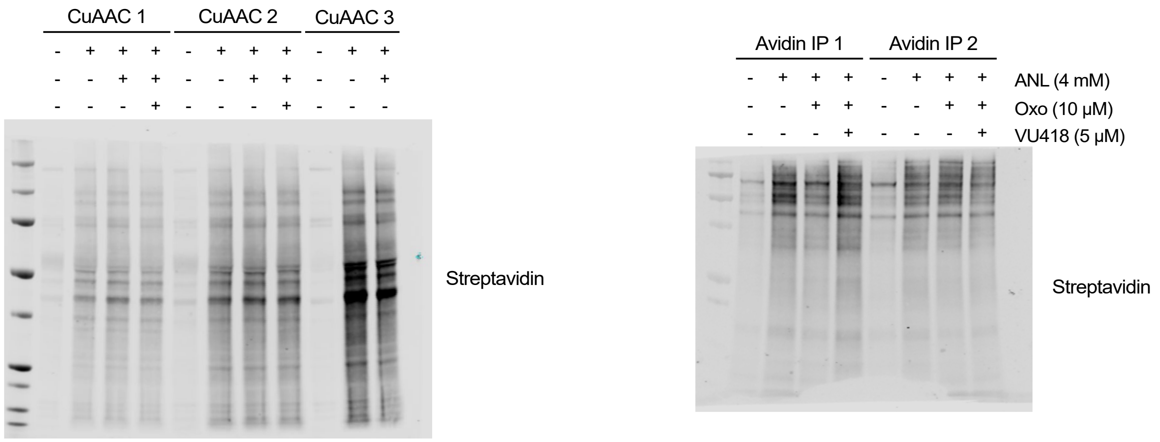

**Uncropped immunoblots related to Suppl. Fig. 6c and 6d**

**Striatum samples**

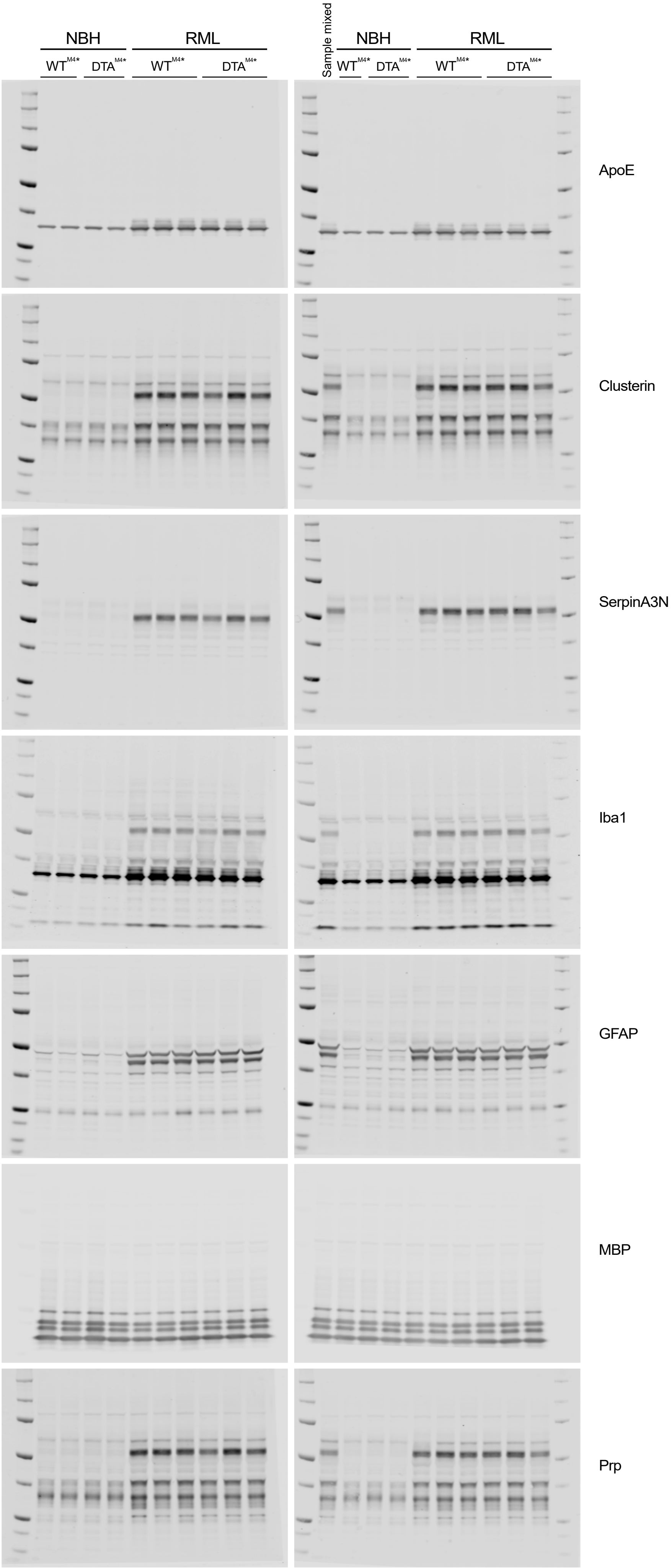

**Uncropped immunoblots related to Suppl. Fig. 6c and 6d**

**Cortex samples**

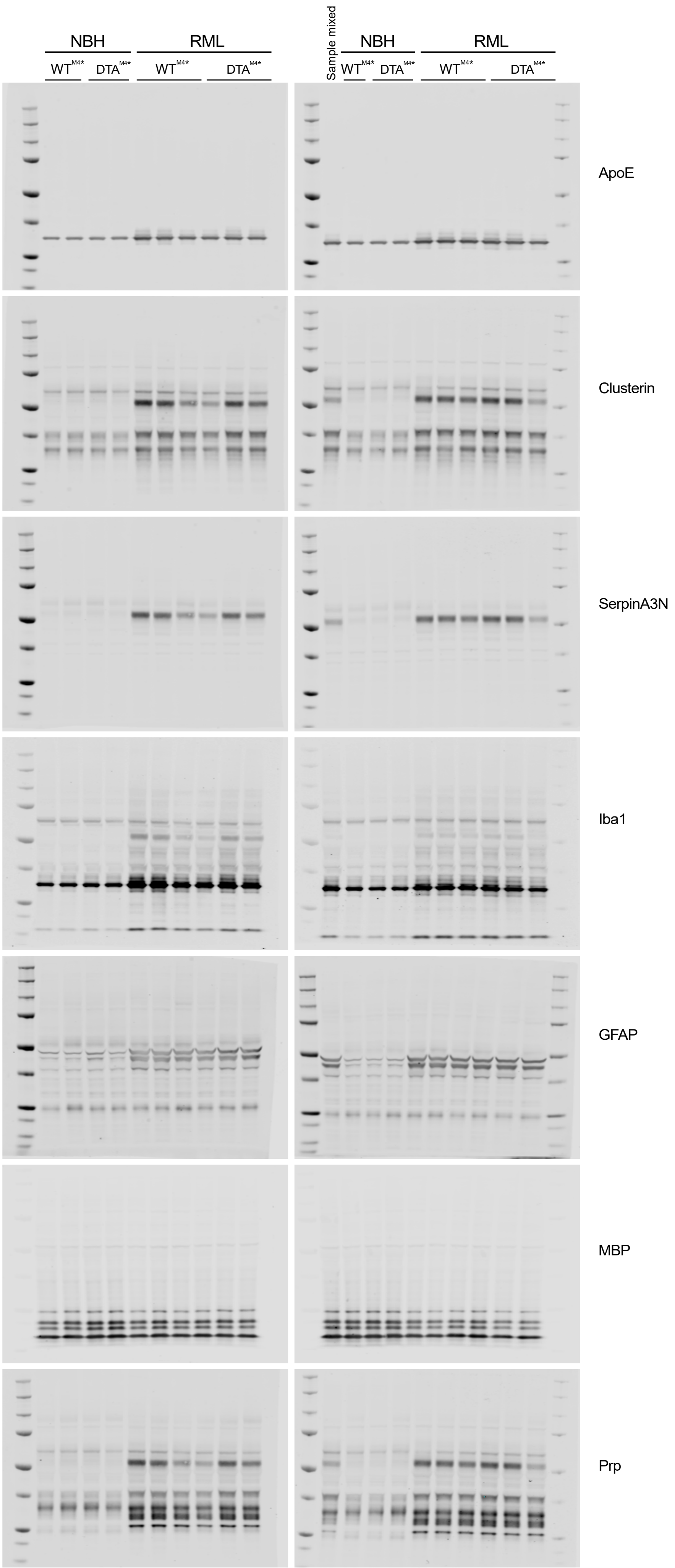

**Uncropped immunoblots related to Suppl. Fig. 6c and 6d**

**Hippocampus samples**

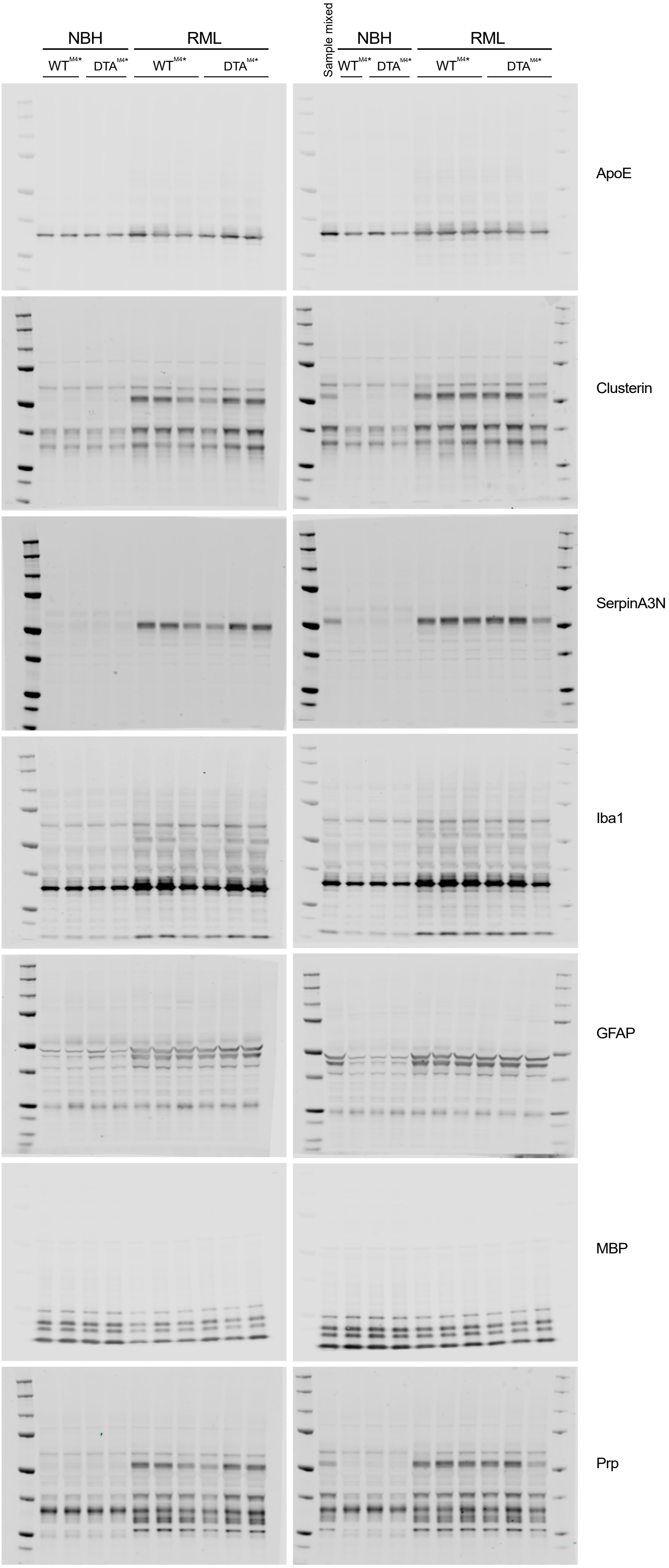

**Uncropped immunoblots related to Suppl. Fig. 6c and 6d**

**Cerebellum samples**

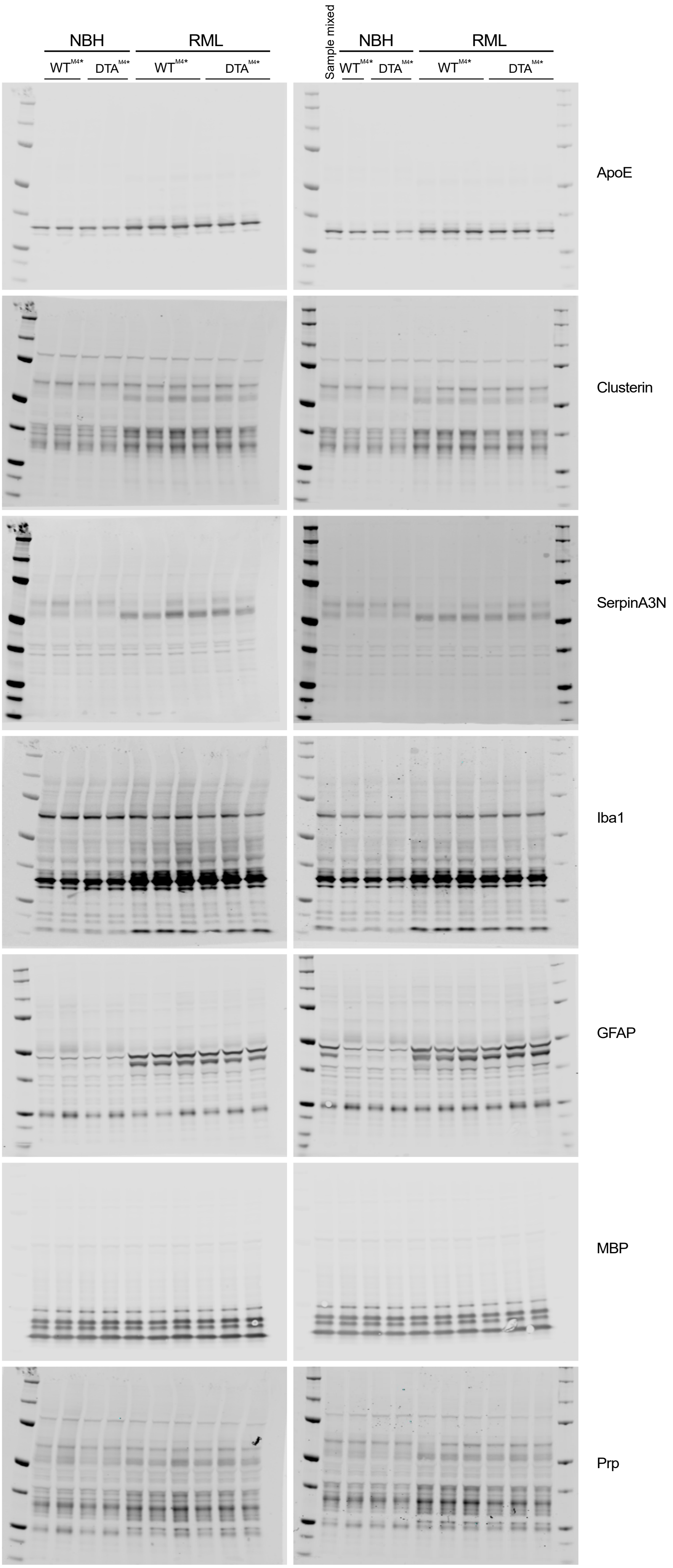
